# AGO2 slicing restrains RTL1 to prevent vascular defects and lethal neonatal muscle failure

**DOI:** 10.1101/2025.04.02.646793

**Authors:** Manish Kumar, Andrea G. Maria, Aurelia Moses, Mahendra Prajapat, Jenna Clements, Moe Kitazawa, Hirosuke Shiura, Kei Fukuda, Joana A. Vidigal

**Affiliations:** Laboratory of Biochemistry and Molecular Biology, National Cancer Institute, The National Institutes of Health, Bethesda, MD, USA; Biophysics Graduate Program, University of Maryland, College Park, MD 20742, USA; Faculty of Life and Environmental Sciences, University of Yamanashi, Yamanashi 400-8510, Japan

## Abstract

Argonaute-mediated RNA slicing is an ancient genome-defense mechanism linked to transposon repression. Although RNAi is no longer a major repeat-control pathway in mammals, AGO2 has retained catalytic activity, and this activity is essential for postnatal viability. The physiological targets that explain this requirement have remained unknown. Here we show that mammalian AGO2 slicing is required to restrain *Rtl1*, an imprinted domesticated retrotransposon whose cleavage is directed by miRNAs from the *miR-433∼127* cluster. Loss of AGO2 catalysis derepresses *Rtl1* and causes endothelial defects and lethal neonatal muscle failure. Mechanistically, RTL1 is an oligomerization-prone retrotransposon-derived protein whose excess expression in muscle induces a non-aggregate proteostasis-stress state characterized by increased proteasome-dependent activity, reduced nascent protein synthesis, and depletion of mitochondrial, calcium-handling, and contractile gene programs. Thus, an essential function of mammalian AGO2 catalysis is to restrain a domesticated transposon-derived protein whose uncontrolled expression compromises endothelial and skeletal muscle function. These findings reveal how an ancestral transposon–Argonaute regulatory interaction has been repurposed to support mammalian development and survival.

## INTRODUCTION

Transposable elements (TEs) are selfish repetitive sequences that pose a persistent threat to genome integrity because they can mobilize to new genomic locations (Bourque, Burns et al. 2018, Wells and Feschotte 2020) which can disrupt essential genes or regulatory elements and thus compromise tissue function and organismal health (Klattenhoff, Bratu et al. 2007, Toth, Pezic et al. 2016, Kazazian and Moran 2017, Rodriguez-Martin, Alvarez et al. 2020, Burns 2022). To limit these risks, organisms have evolved many mechanisms to restrain TE activity (Maksakova, Mager et al. 2008), including RNA interference (RNAi) (Watanabe, Totoki et al. 2008, Sala, Kumar et al. 2023). RNAi is an ancestral RNA-silencing pathway (Fire, Xu et al. 1998) thought to have arisen as a rudimentary innate defense system against exogenous viruses and endogenous repeats (Obbard, Gordon et al. 2009). At the center of this pathway is a catalytic Argonaute (AGO) protein, which uses a bound small-RNA guide to directly cleave, or “slice,” target transcripts with perfect or near-perfect complementarity (Zamore, Tuschl et al. 2000, Becker, Ober-Reynolds et al. 2019). This is an efficient strategy to repress retrotransposons, a class of TEs that mobilize via an RNA intermediate (Wicker, Sabot et al. 2007).

Although Argonaute-mediated cleavage remains a common mechanism for repressing retrotransposons and other repeats in many eukaryotes (Vastenhouw and Plasterk 2004, Matzke and Birchler 2005, Chung, Okamura et al. 2008, Chapman, Taglini et al. 2022), it is no longer a major repeat-control pathway in mammals. Instead, mammalian Argonautes have been largely repurposed for endogenous gene regulation through the microRNA pathway, in which target repression generally does not require slicing (Jonas and Izaurralde 2015, Bartel 2018, Sala, Chandrasekhar et al. 2020). Despite this shift, AGO2, one of the four mammalian Argonaute proteins, has retained catalytic competence through the highly conserved DEDH tetrad in its PIWI domain (Liu, Carmell et al. 2004, Nakanishi, Weinberg et al. 2012). This catalytic activity is essential for postnatal viability: mice homozygous for a point mutation that inactivates the AGO2 catalytic center (*Ago2^D598A/D598A^*) die shortly after birth (Cheloufi, Dos Santos et al. 2010, Jee, Yang et al. 2018). However, the cause of this lethality and the endogenous cleavage target or targets responsible for it have remained unknown. As a result, the physiological gene-regulatory functions of mammalian RNAi remain poorly defined.

To address this question, we re-examined litters from *Ago2^+/D598A^*intercrosses and performed a comprehensive characterization of AGO2 catalytic-mutant animals. We found that homozygous mutants exhibit extensive developmental abnormalities, including systemic vascular and skeletal muscle defects. Conditional genetics revealed that loss of AGO2 catalytic activity in skeletal muscle, but not in the endothelial lineage, is sufficient to recapitulate the neonatal lethality of *Ago2^D598A/D598A^*animals, indicating that muscle dysfunction is the primary cause of death in the absence of AGO2 slicing. Consistent with this, constitutive and skeletal muscle-specific catalytic mutants die shortly after birth with signs of respiratory distress, which we attribute to defects in the major muscles required for breathing. Mechanistically, and befitting Argonaute’s ancestral role in TE restriction, we show that the vascular and muscular defects of AGO2 slicer mutants are caused by derepression of *Rtl1*, a paternally expressed imprinted gene derived from a domesticated retrotransposon. In wild-type animals, *Rtl1* is cleaved by AGO2 through miRNAs from the maternally expressed *miR-433∼127* cluster. In muscle, uncontrolled RTL1 expression caused by either loss of AGO2 slicing competence or loss of the *miR-433∼127* cluster induces a proteostasis-stress program and disrupts mitochondrial, calcium-handling, and contractile gene programs. Together, our work identifies *Rtl1* as an essential endogenous mRNA cleavage target of AGO2 and suggests that a domesticated retrotransposon retained an ancestral-like dependence on catalytic Argonaute regulation, contributing to the evolutionary retention of AGO2 slicing activity in mammals.

## RESULTS

### Vascular and muscular abnormalities in the absence of AGO2 slicing competence

To define the phenotypic requirements for AGO2 catalytic activity in mice, we analyzed animals in which this activity is selectively disrupted through a point mutation in the conserved catalytic domain of the protein (*Ago2^D598A^*) (Cheloufi, Dos Santos et al. 2010). As previously reported, homozygous mutants were recovered at the expected Mendelian ratio at embryonic day (E) 18.5 but died within hours of birth with evidence of anemia, a previously described defect in these mutants (Cheloufi, Dos Santos et al. 2010, Jee, Yang et al. 2018). Although severe (Cheloufi, Dos Santos et al. 2010), anemia is not the cause of *Ago2^D598A/D598A^* lethality since conditional loss of catalytic competence in the hematopoietic system recapitulates the erythroid defect but is compatible with life (Jee, Yang et al. 2018). We therefore performed a broader phenotypic characterization of embryos lacking AGO2 slicing activity.

First, we observed a dose-dependent increase in placental weight (**Supplementary Figure 1A**) with no significant change in embryo weight (**Supplementary Figure 1B**), leading to a significantly reduced placental efficiency (**Supplementary Figure 1C**). Higher placental weight correlated with increased levels of phosphorylated histone H3 consistent with increased cell proliferation in this organ (**Supplementary Figure 1D, 1E**). Placental phenotypes were recapitulated in *Ago2^flx/D598A^;Sox2-Cre^+/-^*animals (**Supplementary Figure 1F-H, Supplementary Figure 2**) showing they originate from a dysfunction in epiblast-derived cells and not from the trophectoderm lineage. Since epiblast-derived cells in the placenta give rise to endothelial cells that line the fetal vessels of that organ (Rossant and Cross 2001), we tested if loss of AGO2 catalytic competence impacts placental vascular development. We found that both *Ago2^D598A/D598A^*and *Ago2^flx/D598A^;Sox2-Cre^+/-^* animals had abnormally patterned vessels in the placentas, which were significantly enlarged compared to controls (**Figure 1A**, **Supplementary Figure 1I**). These vascular defects extended to the fetus as well, which at E18.5 also showed evidence of vessel dilation (**Figure 1B**). Moreover, catalytic mutant animals frequently presented with internal hemorrhages at birth (**Figure 1C**), in line with previous work (Cheloufi, Dos Santos et al. 2010). This could reflect impaired endothelial barrier function, which in extreme cases causes internal bleeding.

**Fig. 1.**
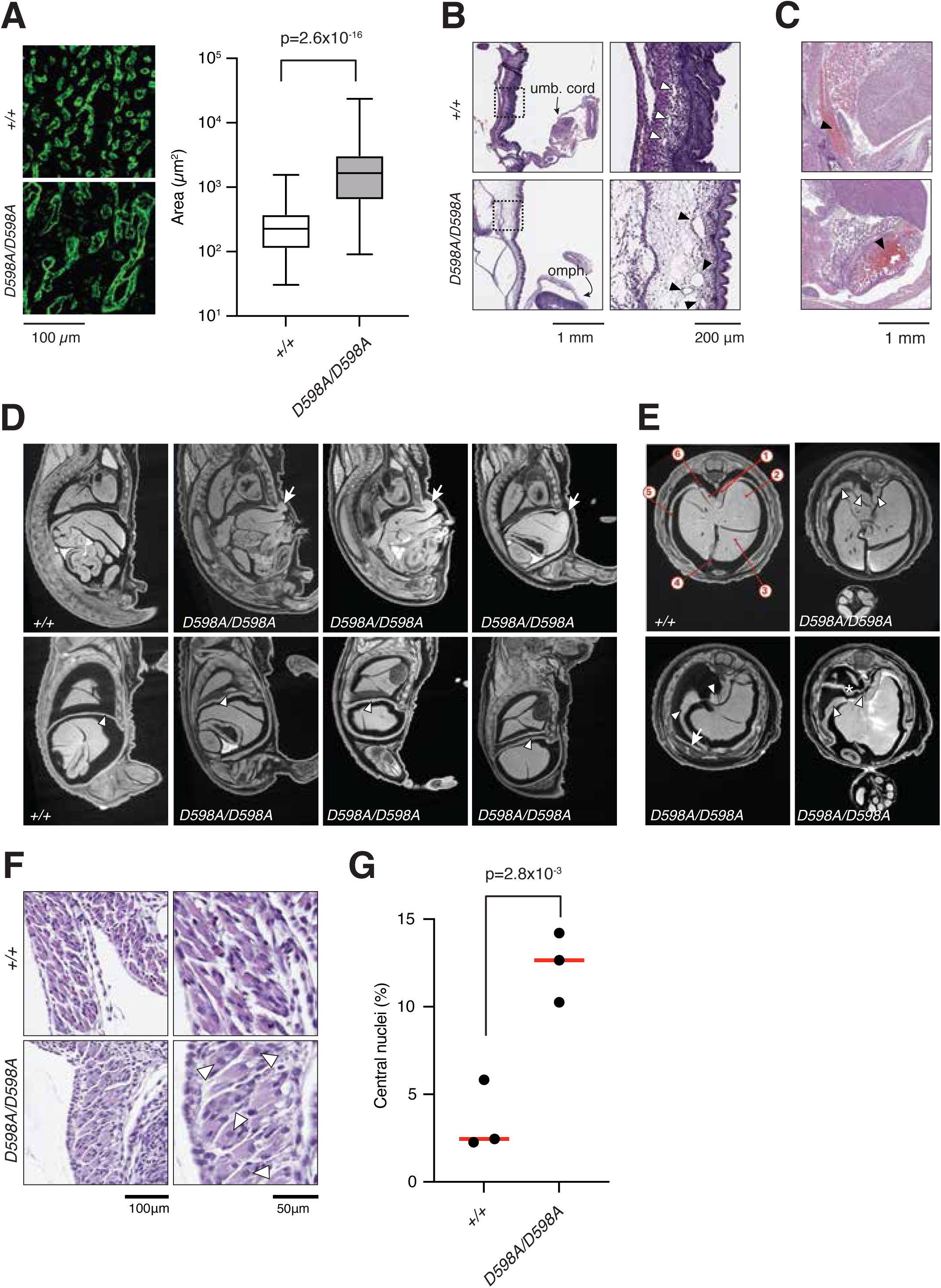
Extensive developmental defects in the absence of AGO2 cleavage competence. **(A)** Left, immunofluorescence staining of mouse placentas with an antibody against the CD31 endothelial cell marker. Right, quantification of vessel area in wild-type and mutant placentas. p-values were calculated using a two-tailed t-test. **(B)** Hematoxylin-Eosin-stained sagittal sections of E18.5 embryos, showing enlarged vessels in mutant (bottom, closed arrowheads) compared to wild-type (top, open arrowheads) animals. **(C)** Hematoxylin-Eosin-stained sagittal sections showing examples of abdominal hemorrhages (arrowhead) in *Ago2* catalytic mutants. **(D)** Micro-CT sagittal sections of wild-type (+/+) and catalytic mutant (D598A/D598A) animals at E18.5. Arrows point to liver deformation in mutants; arrowheads highlight the diaphragm. **(E)** Micro-CT transverse sections of wild-type (+/+) and catalytic mutant (D598A/D598A) animals at E18.5. Note the thickened transversus thoracis muscle (arrow) and pleuropericardial membrane (asterisk). Arrowheads highlight diaphragm muscle and crura which are also significantly thickened and misshapen compared to controls. Labels: 1, right and left crux of diaphragm muscle; 2, left lobe of liver; 3, left middle lobe of liver; 4, Falciform ligament, 5, diaphragm; 6, diaphragm. **(F)** Hematoxylin-Eosin-stained sections of diaphragm muscle at P0. Arrowheads point to muscle fibers with centrally located nuclei. **(G)** Quantification of the percentage of myofibers containing centrally located nuclei. Each dot represents an animal. p-values were calculated using a two-tailed t-test.

Second, *Ago2^D598A/D598A^* animals presented with numerous abdominal malformations. The most prominent macroscopic defect at E18.5 was omphalocele, an abdominal wall closure defect in which the intestine protrudes outside the body cavity but remains inside a membranous sac (**Supplementary Figure 2A-D**). This defect correlated with the presence of reduced visceral cavity size in these animals and was largely absent by birth, suggesting a delay in resolving the physiological umbilical hernia that is naturally present in wild-type animals from E12.5-E16.5. μCT analysis also showed that all mutant animals examined (n=9) displayed misshapen liver lobes, likely secondary to their partial extrusion from the visceral cavity (**Figure 1D**, top). The scans also revealed abnormally thick body-wall musculature in catalytic mutant embryos, particularly evident in the transversus thoracis muscle (**Figure 1E**, arrow). An abnormally thick pleuropericardial membrane was also evident in some embryos (**Figure 1E**, asterisk).

Alongside these minor defects in cavity membranes and abdominal muscle, we observed major defects in the diaphragms of all mutants. Specifically, not only were the mutant diaphragm muscle and the diaphragm crura thicker than those of controls (**Figure 1D**, bottom; **Figure 1E**, arrowheads; **Supplementary Figure 2E**) but in all cases the diaphragm displayed an irregular, jagged morphology in contrast to the smooth shape of wild-type diaphragms. Hematoxylin and eosin staining of embryo sections further revealed structural abnormalities in mutant skeletal muscle, particularly a markedly increased proportion of centrally nucleated myofibers (**Figure 1F**, **1G**). Because centrally located nuclei can be associated with regeneration or delayed maturation, we examined expression of embryonic and perinatal myosin heavy-chain genes. *Myh3* was not induced in *Ago2^D598A/D598A^*muscles, and *Myh8* was modestly reduced (**Supplementary Figure 2G**). Thus, central nucleation is not explained by persistence or reactivation of an embryonic/perinatal myosin program, but instead likely reflects impaired myofiber architecture or nuclear positioning.

Finally, we examined whether catalytic mutants displayed skeletal patterning defects because *Hoxb8* is one of the few known cleavage targets of AGO2 in the developing embryo (Yekta, Shih et al. 2004). Despite the importance of HOXB8 for normal vertebral development (van den Akker, Reijnen et al. 1999), the posterior homeotic transformations caused by *Hoxb8* overexpression (Charite, de Graaff et al. 1994), and the conservation of the *miR-196*/*Hoxb8* cleavage interaction across species (Yekta, Shih et al. 2004, He, Yan et al. 2011), we found no skeletal patterning defects in *Ago2^D598A/D598A^* animals (**Supplementary Figure 3**). We did however notice that all *Ago2^D598A/D598A^*embryos displayed some degree of growth retardation, pronounced spinal curvatures, and shorter bones compared to controls, although these are not reflected in the weights.

Together, these results represent a comprehensive characterization of mice lacking *Ago2* catalytic competence. Our data show that in addition to the well-documented block in erythrocyte development (Cheloufi, Dos Santos et al. 2010, Jee, Yang et al. 2018), these animals display vascular, placental, abdominal wall, and skeletal muscle abnormalities, revealing a broad requirement for AGO2-mediated slicing in mammalian development. Importantly, the two major defects we document here—in vasculature and respiratory muscles—affect systems whose disruption can compromise perinatal viability (Joureau, de Winter et al. 2017, Perez-Garcia, Fineberg et al. 2018), and thus may contribute to the lethality of *Ago2^D598A/D598A^*animals.

### Transcriptomic impact of AGO2 slicing activity

To understand the broader impact of AGO2 slicing during mouse development and to identify the cleavage target underlying the defects above we sequenced both placenta and diaphragm tissue from *Ago2^D598A/D598A^* embryos just before birth. Loss of AGO2 slicing resulted in numerous dysregulated genes in both tissues (**Figure 2A, 2B**), with a notable imbalance towards upregulated genes in mutant placentas. This likely reflects altered placental cell composition in the absence of AGO2 catalytic competence, since gene ontology terms related to vascular development and angiogenesis were significantly enriched among upregulated genes (**Supplementary Figure 4A**), together with gene signatures of vascular and lymphatic endothelial cell-types (Marsh and Blelloch 2020) (**Supplementary Figure 4B**).

**Fig. 2.**
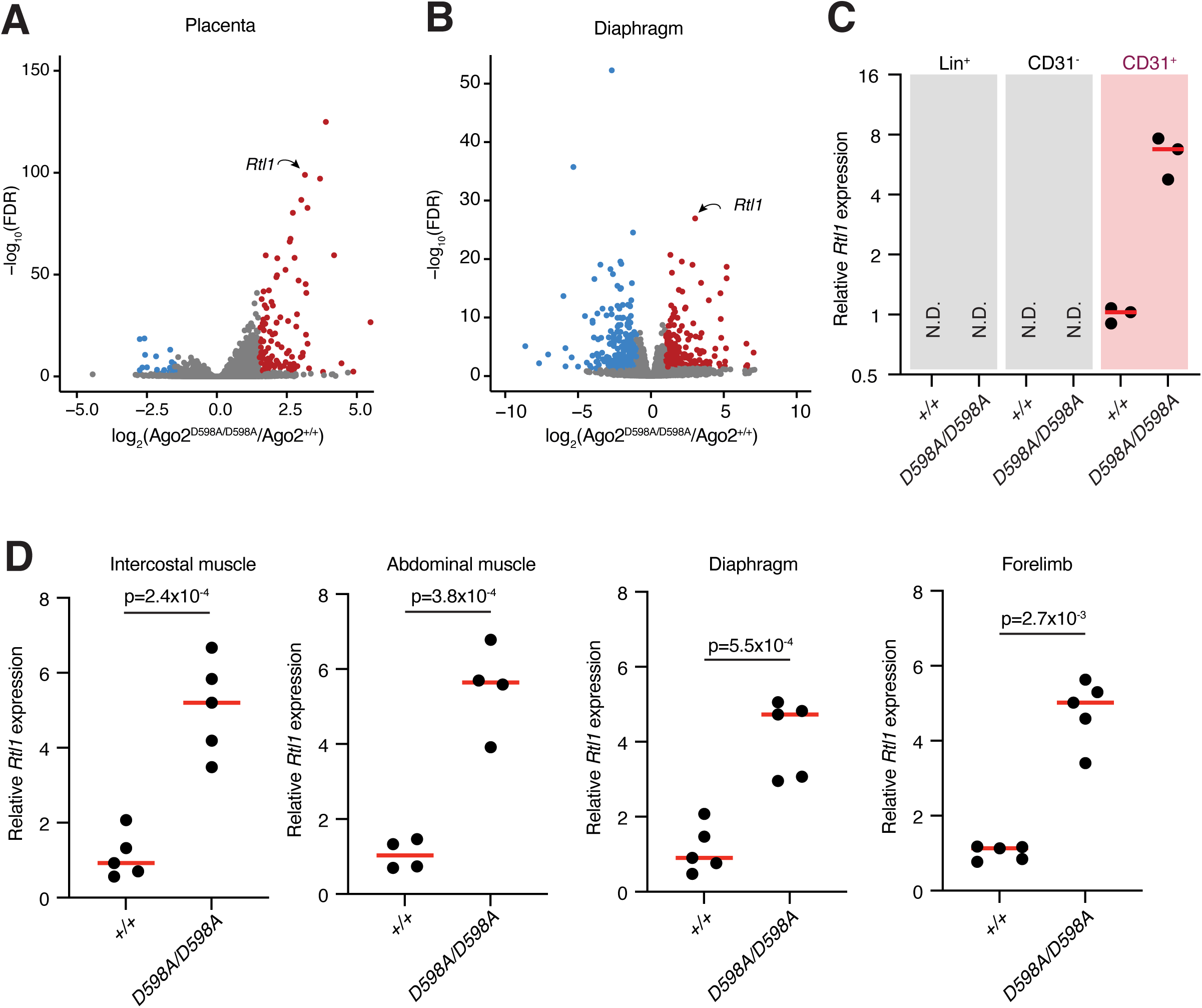
Transcriptional changes downstream of AGO2 slicing loss. **(A)** Volcano plot showing genes significantly dysregulated in *Ago2* mutant placentas (FDR < 0.05 and |log2(fold change)| > 1). **(B)** Volcano plot showing genes significantly dysregulated in *Ago2* mutant diaphragms (FDR < 0.05 and |log2(fold change)| > 1). **(C)** *Rtl1* is expressed and upregulated in endothelial (CD31+) cells isolated from placentas N.D., not detected. **(D)** *Rtl1* is expressed and upregulated in skeletal muscle from various origins. Individual dots in (C) and (D) represent different animals. p-values were calculated using a two-tailed t-test.

The anemia phenotype previously reported in *Ago2^D598A/D598A^* animals (Cheloufi, Dos Santos et al. 2010, Jee, Yang et al. 2018) is caused by a block in the maturation of *miR-451* and *miR-486*, two erythrocyte-specific miRNAs whose processing requires AGO2 catalysis (Cheloufi, Dos Santos et al. 2010, Jee, Yang et al. 2018). These are currently the only two RNA cleavage events whose disruption have been shown to cause phenotypic defects in *Ago2^D598A/D598A^*mice. We therefore considered the possibility that the phenotypes and transcriptional changes we observed in the absence of AGO2 slicing competence might also be secondary to the deregulation of another miRNA whose biogenesis is AGO2-dependent. We found no evidence that this was the case. First, only a few mature miRNAs differed between wild-type and mutant animals in our dataset, totaling 12 dysregulated miRNAs (9 upregulated, 3 downregulated; **Supplementary Figure 5A**, **Supplementary Table 1**). All these miRNAs except for *miR-451* arise from characteristically structured hairpin precursors, suggesting that they mature through a typical biogenesis pathway that does not require AGO2 slicing. Second, predicted targets of these miRNAs were not preferentially dysregulated in *Ago2^D598A/D598A^*samples (**Supplementary Figure 5B-C**), suggesting that the most miRNA changes are indirect consequences of the large transcriptomic differences we have observed rather than their direct drivers.

Alternatively, we hypothesized that some miRNAs could direct AGO2 to cleave mRNAs whose consequent upregulation would cause the developmental defects described here. To test this hypothesis, we computationally searched among upregulated transcripts for the presence of sequences with perfect complementarity to miRNAs with at least 30 reads per million (RPM) in our datasets (**Supplementary Table 2**). A single transcript stood out from this analysis, *Rtl1. Rtl1* was significantly upregulated in our mutants (**Figure 2A-D, Supplementary Figure 4C-F**) and contained 10 perfectly complementary sites to well-expressed miRNAs (**Supplementary Table 3**), suggesting that it could be a direct cleavage target of AGO2 *in vivo*. Importantly, *Rtl1* has been shown to have roles in both placental fetal endothelial (Sekita, Wagatsuma et al. 2008) and muscle (Kitazawa, Hayashi et al. 2020) differentiation, with its overexpression phenocopying most of the abnormalities we observed in *Ago2* catalytic mutants. These include enlarged placental fetal vessels and placentomegaly without changes in fetal weight (Sekita, Wagatsuma et al. 2008), structural abnormalities affecting diaphragm, intercostal, and abdominal skeletal muscles, including fibers with centrally located nuclei (Kitazawa, Hayashi et al. 2020); and perinatal lethality (Sekita, Wagatsuma et al. 2008, Kitazawa, Hayashi et al. 2020). Notably, *Rtl1* was significantly upregulated in both extra-embryonic and embryonic tissues, including in the endothelium (defined as CD31^+^ cells) (**Figure 2C**; **Supplementary Figure 4F**) and in embryonic muscles of the diaphragm, intercostal region, abdomen, and forelimbs (**Figure 2D**). Based on these data, we hypothesized that *Rtl1* is the key AGO2 cleavage target whose upregulation leads to both the developmental defects we have identified here and to the previously unexplained neonatal lethality of catalytic mutant mice.

### *Rtl1*, a domesticated transposon, is a cleavage target of AGO2

*Rtl1* (Retrotransposon-like protein 1; also known as *Peg11*, paternally expressed gene 11) is an imprinted domesticated retrotransposon that is expressed solely from the paternal chromosome (**Figure 3A**) (Charlier, Segers et al. 2001, Cavaille, Seitz et al. 2002). The maternal chromosome expresses instead an imprinted noncoding RNA that overlaps the *Rtl1* locus and serves as the primary transcript for a cluster of 6 miRNAs (*miR-136*, *miR-432*, *miR-434*, *miR-127*, *miR-433*, *miR-431*) (**Figure 3A**) (Seitz, Youngson et al. 2003), five of which are above the 30 RPM threshold in our dataset (*miR-136*, *miR-434*, *miR-127*, *miR-433*, *miR-431*). Because these miRNAs are transcribed in an antisense manner relative to *Rtl1*, their sequences are perfectly complementary to that gene (**Figure 3B**, **Supplementary Table 3**) and can trigger AGO2-mediated cleavage of *Rtl1* at the position opposite to nucleotides 10 and 11 of the small RNA sequence (Davis, Caiment et al. 2005). We looked for evidence of such cleavage products in total RNA extracted from wild-type animals using a modified 5’RACE protocol. We detected cleavage products for all five well-expressed miRNAs from this cluster, and in two instances for passenger strands as well, for a total of 7 confirmed cleavage sites on the *Rtl1* transcript (**Figure 3C**). These cleavage products were absent from RNAs extracted from *Ago2^D598A/D598A^* samples indicating that they are a direct result of AGO2’s slicing activity (**Figure 3C**). Importantly, sequencing of these 5’RACE amplicons confirmed that the 5’adaptor had ligated to a 5’phosphate group at the predicted cleavage position within the target sequence in most molecules (**Figure 3B, 3D**). Thus, our data confirm that *Rtl1* is a true cleavage target of AGO2 *in vivo* and identify at least 7 physiological AGO2 cleavage events at this transcript during mouse embryogenesis that are disrupted in *Ago2* catalytic mutant animals.

**Fig. 3.**
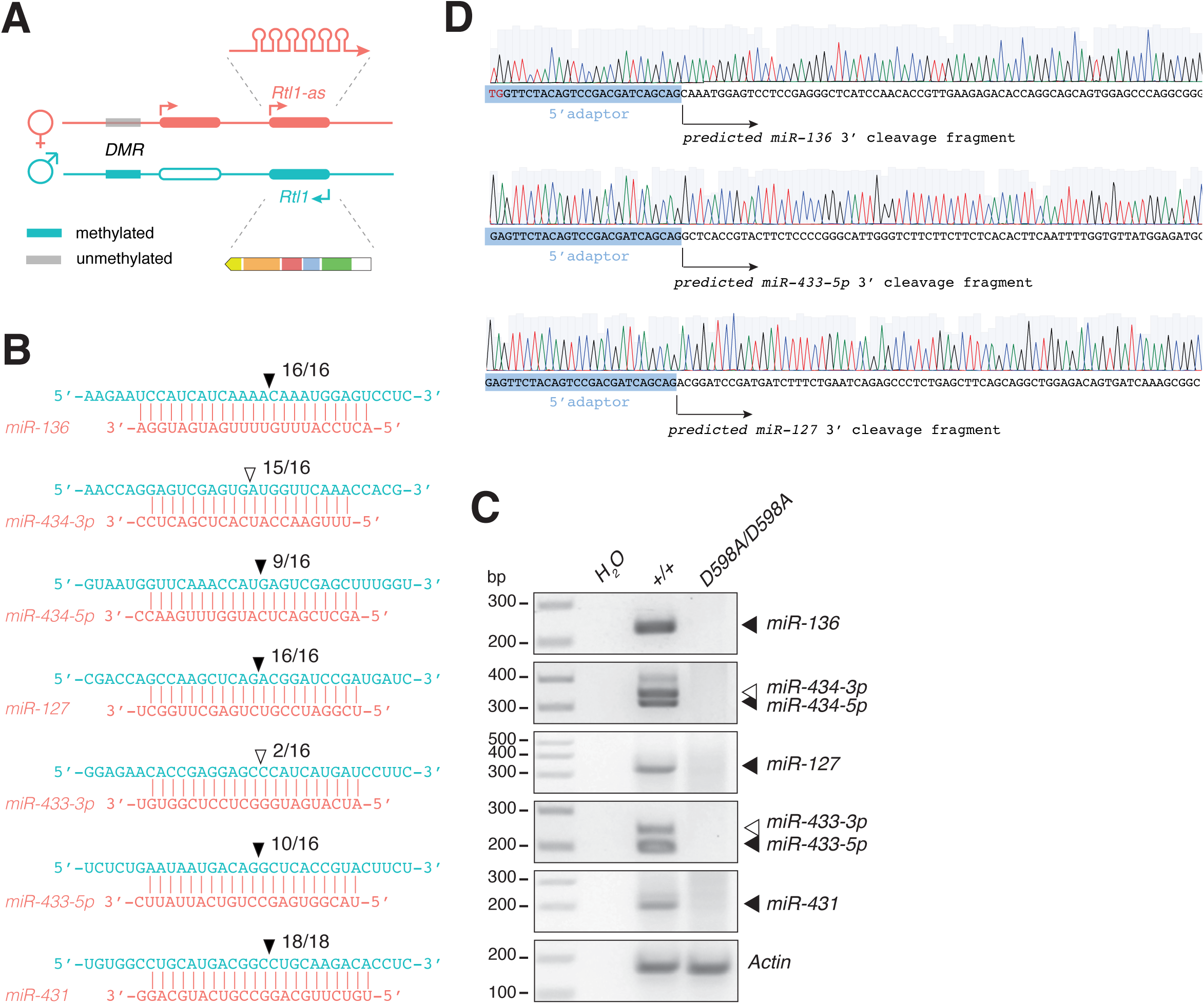
*Rtl1* is a cleavage target of AGO2 during embryogenesis. **(A)** Schematic representation of the *Rtl1* imprinted locus. The differentially methylated region (DMR) of this locus, which is methylated only in the paternal allele leads to imprinted expression of multiple genes including *Rtl1* and *miR-433∼127* whose expressions overlap in an antisense manner. *miR-433∼127* is processed into 6 distinct miRNAs. **(B)** Representation of the pairing between miRNAs of the *miR-136*∼*miR-431* cluster (orange) and the *Rtl1* transcript (blue). The position of the predicted cleavage site is indicated, along with the number of times that cleavage site was sequenced. **(C)** Identification of AGO2 cleavage products in wild-type (+/+) but not catalytic mutant (D598A/D598A) samples. The identity of the miRNA triggering the cleavage event is shown on the right. Closed and open arrowheads denote mature and star strands respectively. **(D)** Representative Sanger sequencing traces showing that 5′RACE amplicons correspond to the predicted AGO2 cleavage products.

### Maternal deletion of the *Rtl1* locus recapitulates the transcriptome of *Ago2* slicing mutants

To determine whether derepression of *Rtl1* accounts for the phenotypes observed in the placenta and diaphragm of *Ago2* slicer mutants, we collected both tissues at E18.5 from embryos carrying a maternally inherited deletion of the *Rtl1* locus (*Rtl1^ΔMat/+^*) and performed RNA sequencing. This line, which had been backcrossed to the C57BL/6 background for ten generations, was previously shown to exhibit developmental phenotypes that overlap those observed here in *Ago2^D598A/D598^* embryos (Sekita, Wagatsuma et al. 2008, Kitazawa, Hayashi et al. 2020). Due to imprinting, *Rtl1^ΔMat/+^* animals retain the expression of *Rtl1* but not of the antisense polycistronic miRNA cluster that targets it for slicing (**Figure 4A**). In line with this, loss of the maternal locus resulted in significant upregulation of the *Rtl1* transcript in both tissues and these changes were comparable to those observed in the absence of AGO2 slicing (**Figure 4B, 4C**). Thus, *Rtl1^ΔMat/+^* embryos provide an independent genetic model in which to test the extent to which transcriptional changes caused by loss of AGO2 catalytic competence are recapitulated by *Rtl1* derepression.

**Fig. 4.**
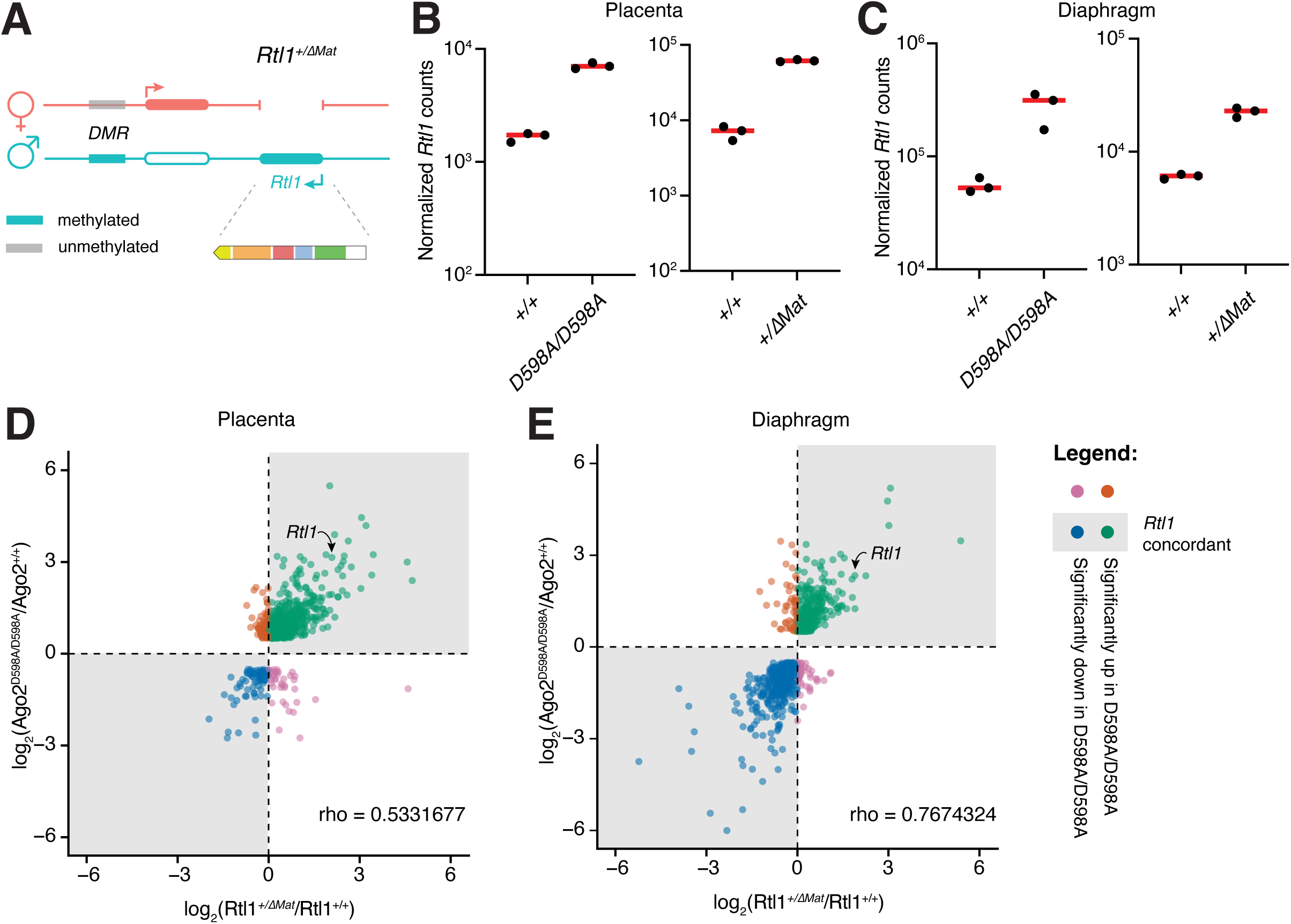
Deletion of the *miR-433∼127* cluster recapitulates transcriptomic changes of Ago2 catalytic mutants. **(A)** Schematic representation of the *Rtl1^ΔMat/+^* allele. **(B)** Normalized *Rtl1* counts in *Ago2^D598A/D598A^* (left) and *Rtl1^ΔMat/+^* (right) placentas. **(C)** Normalized *Rtl1* counts in *Ago2^D598A/D598A^*(left) and *Rtl1^ΔMat/+^*(right) diaphragms. **(D**, **E)** For each tissue, genes significantly differentially expressed in AGO2 catalytic mutants were plotted according to their log2 fold change in *Rtl1^ΔMat/+^* and *Ago2^D598A/D598A^*mutants. Genes in the upper-right and lower-left quadrants change in the same direction in both mutants and are interpreted as transcriptionally concordant. Spearman correlation coefficients are shown.

To do so, we compared the transcriptional consequences of *Ago2* slicing loss with those produced by *Rtl1^ΔMat/+^*. For each tissue, we found that significant transcriptional changes observed in *Ago2^D598A/D59AA8^*embryos were largely recapitulated in *Rtl1^ΔMat/+^* animals. Specifically, in placenta, *Ago2*-significant genes showed a positive correlation with the *Rtl1^ΔMat/+^*signature (Spearman rank correlation of ρ = 0.533) with 84.3% of changes being directionally concordant between the two mutants (**Figure 4D**). Concordance was stronger in diaphragm which showed a higher correlation between datasets (Spearman rank correlation of ρ = 0.767), with 87.6% of the genes differentially expressed in *Ago2^D598A/D59AA8^* muscle changing in the same direction when the miRNA cluster was deleted (**Figure 4E**). The weaker correlation we observed in placenta is consistent with the greater cellular complexity of this tissue and with the possibility that a subset of placental expression changes reflects altered tissue composition or other indirect effects, rather than direct transcriptional consequences of *Rtl1* upregulation alone.

Taken together, our data show that direct upregulation of *Rtl1* through genetic deletion of the miRNAs that target it for slicing results in genome-wide transcriptional changes that largely overlap those observed in *Ago2* catalytic-mutant samples. This supports a model in which *Rtl1* is the major physiologically relevant AGO2 slicing target in both the placenta and the diaphragm, and in which *Rtl1* upregulation accounts for most of the transcriptomic and phenotypic abnormalities observed in *Ago2^D598A/D59A8A^* embryos.

### Endothelial cell defects do not cause lethality of *Ago2* slicing mutants

The two major defects documented here—in endothelial and respiratory muscle development—affect systems whose disruption can compromise perinatal viability (Joureau, de Winter et al. 2017, Perez-Garcia, Fineberg et al. 2018). To test whether either defect underlies the lethality of *Ago2^D598A/D598A^* animals, we generated animals that conditionally lose AGO2 slicing competence in either endothelial-lineage cells or skeletal muscle.

First, to determine whether vascular defects underlie the neonatal lethality in *Ago2* catalytic mutant animals, we crossed *Ago2^+/D598A^; Tie2-Cre^+/-^* males to *Ago2^flx/flx^* females, leading to Cre expression in the endothelial lineage of the resulting embryos and specific recombination of the floxed allele in the endothelial cells of Tie2-Cre positive animals (**Supplemental Figure 6A**). This resulted in a significant increase in *Rtl1* expression in CD31^+^ endothelial cells (**Figure 5A**). Conditional loss of *Ago2* catalytic competence in these mice also recapitulated the vascular defects of constitutive mutants, reflected in a significant enlargement of vessels (**Figure 5B, 5C**). This confirms the vascular defects are caused by a dysfunction of the embryonic endothelium and that they are largely cell autonomous. Yet, despite recapitulating *Rtl1* overexpression and endothelial dysfunction, we observed no impact on animal viability either at birth (**Figure 5D**) or at weaning (**Figure 5E**). Thus, loss of *Ago2* catalytic competence in the *Tie2^+^*-lineage is fully compatible with early postnatal survival, and the disruption of the normal endothelial function is not sufficient to explain the neonatal lethality of *Ago2^D598A/D598A^*constitutive-mutant animals.

**Fig. 5.**
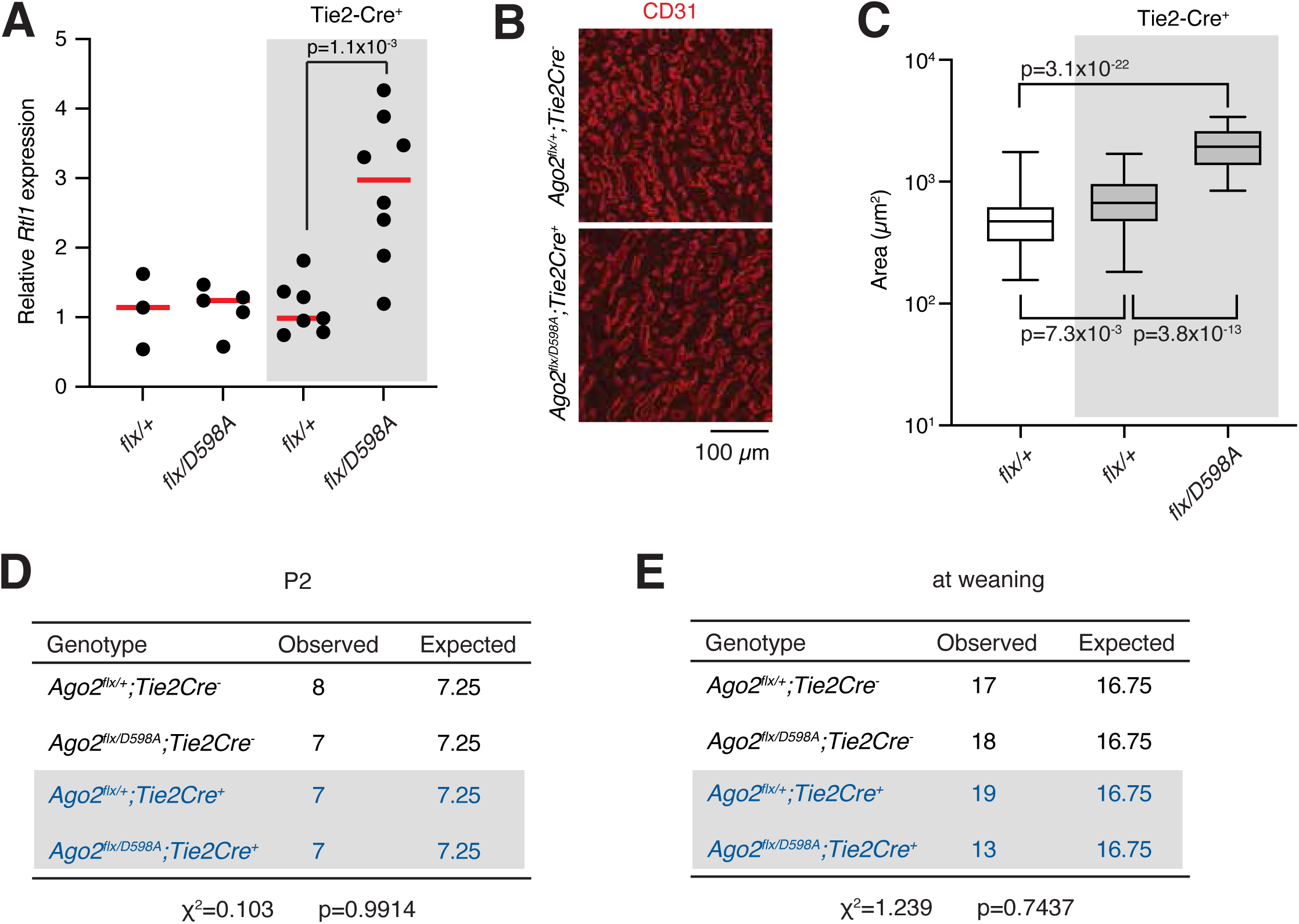
Endothelial-lineage loss of AGO2 slicing recapitulates vascular defects but is compatible with postnatal viability. **(A)** Relative *Rtl1* expression levels in indicated genotypes. Each dot represents an animal. p-values were calculated using a two-tailed t-test. **(B)** immunofluorescence staining of mouse placentas with an antibody against the CD31 endothelial cell marker. **(C)** quantification of vessel area in wild-type and mutant placentas. p-values were calculated using a two-tailed t-test. **(D**, **E)** Genotyping of litters after birth (P2) or at weaning showing conditional loss of AGO2 catalytic competence in the Tie2-lineage is compatible with animal viability. P value was calculated with the Chi-square test.

### Skeletal muscle *Rtl1* derepression causes postnatal lethality in *Ago2* catalytic mutants

To determine whether AGO2 slicing activity in skeletal muscle is required for postnatal viability, we performed an analogous conditional genetic cross. Specifically, we crossed *Ago2^+/D598A^; Acta1-Cre^+/−^* mice to *Ago2^flx/flx^* mice, generating Cre-positive offspring in which the floxed *Ago2* allele is deleted specifically in skeletal muscle. Consistent with efficient recombination of this allele (**Supplementary Figure 6B**), *Rtl1* transcript was significantly upregulated in intercostal (**Figure 6A**), abdominal (**Figure 6B**), and diaphragm muscles (**Figure 6C**) of *Ago2^flx/D598A^; Acta1-Cre^+/−^* animals compared with *Ago2^flx/+^; Acta1-Cre^+/−^* controls. Moreover, this upregulation was comparable to that observed in *Ago2^D598A/D598A^* constitutive mutants (**Figure 2D**).

**Fig. 6.**
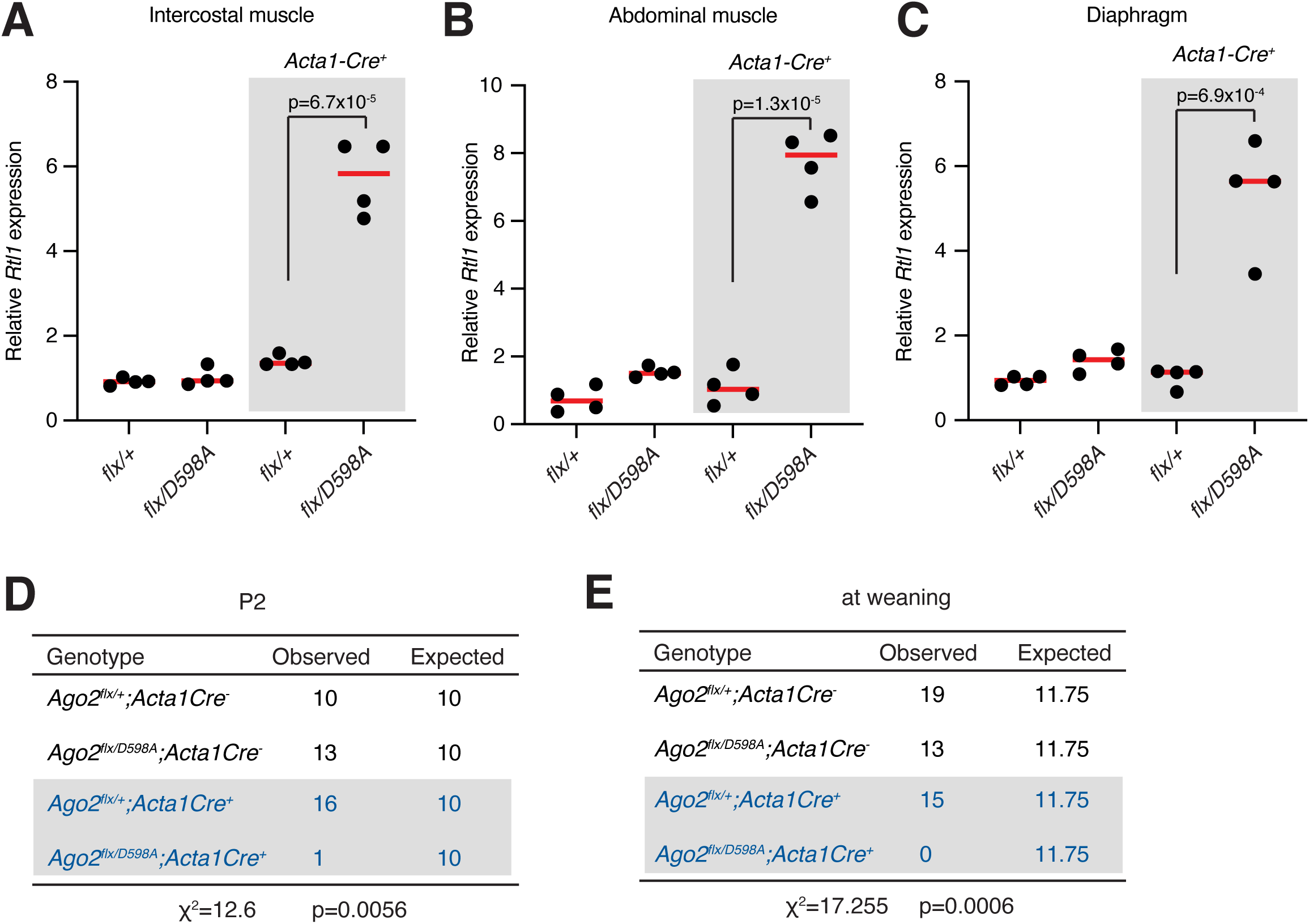
Skeletal muscle loss of AGO2 slicing causes Rtl1 derepression and postnatal lethality. **(A, B, C)** Relative *Rtl1* expression levels in intercostal (A), abdominal (B), and diaphragm (C) muscles in indicated genotypes. Each dot represents an animal. p-values were calculated using a two-tailed t-test. **(D**, **E)** Genotyping of litters after birth (P2) or at weaning showing conditional loss of AGO2 catalytic competence in the Acta1+ skeletal muscle lineage results in postnatal lethality. P value was calculated with the Chi-square test.

Despite efficient deletion of the floxed allele in skeletal muscle, most *Ago2^D598A/flx^*; *Acta1-Cre^+/-^* embryos lacked omphalocele at E18.5 (**Supplementary Figure 6C**), a phenotype with almost complete penetrance in both *Ago2^D598A/D598A^*and *Ago2^D598A/flx^*; *Sox2-Cre^+/−^* animals (**Supplementary Figure 2C-D**). This indicates that the abdominal closure defect is not primarily caused by loss of AGO2 catalytic activity in differentiated Acta1-positive myofibers. Instead, omphalocele in constitutive mutants likely reflects AGO2 catalytic impairment before Acta1-positive myofiber differentiation, in earlier abdominal wall progenitors, or in non-muscle abdominal wall tissues, as described in other mouse models (Nichol, Corliss et al. 2011, Takahashi, Tamura et al. 2018).

We next assessed the impact of loss of Ago2 slicing on animal viability by monitoring animal genotypes after birth and at weaning. Although all genotypes were present at the expected Mendelian ratios immediately before birth (E18.5), only one *Ago2^flx/D598A^; Acta1-Cre^+/−^* animal was recovered among 40 animals genotyped after delivery (**Figure 6D**). No *Ago2^flx/D598A^; Acta1-Cre^+/−^* animals were present among 47 animals that survived to weaning (**Figure 6E**). Thus, in contrast to our findings in the endothelium, loss of AGO2 catalytic activity in skeletal muscle is incompatible with postnatal viability.

Given that loss of AGO2 catalytic competence affects muscles required for breathing—an activity essential for postnatal survival—and that catalytic mutants die shortly after birth with signs of respiratory distress, these data support a model in which lethality of *Ago2^D598A/D598A^* animals results from respiratory failure secondary to muscle abnormalities caused by *Rtl1* overexpression.

### Upregulation of Rtl1 drives proteostasis stress in the muscle of *Ago2* slicing mutants

To understand how *Rtl1* upregulation causes lethal muscle failure in *Ago2* catalytic mutants we performed gene-set enrichment analysis on RNA sequencing datasets from *Ago2^D598A/D598A^* and *Rtl1^ΔMat/+^* diaphragms. Consistent with the strong overlap in gene-level transcriptomic changes between these mutants (**Figure 4E**), the two models showed a highly concordant pathway signature (**Figure 7A**). Among GO terms significantly enriched in both datasets, 320 of 324 showed the same direction of enrichment, indicating a shared transcriptional response. Positively enriched programs were dominated by protein folding/refolding, unfolded protein binding, protein-folding chaperone complexes, and ubiquitin-proteasome regulation. These terms were driven by upregulation of heat-shock foldases and co-chaperones (*Hspa1a*, *Hspa1b*, *Hsph1*, *Hsp90aa1*, *Hsp90ab1*, *Hspa8*, *Dnaja4*, *Dnajb1*, *Dnajb2*, *Stip1*, and *Ahsa1*) together with a small HSP/holdase and BAG3-associated module (*Cryab*, *Hspb1*, *Hspb8*, and *Bag3*) in both mouse models (**Figure 7B**). Thus, *Rtl1* derepression is associated with activation of a shared protein-quality-control program in developing diaphragm muscle.

**Fig. 7.**
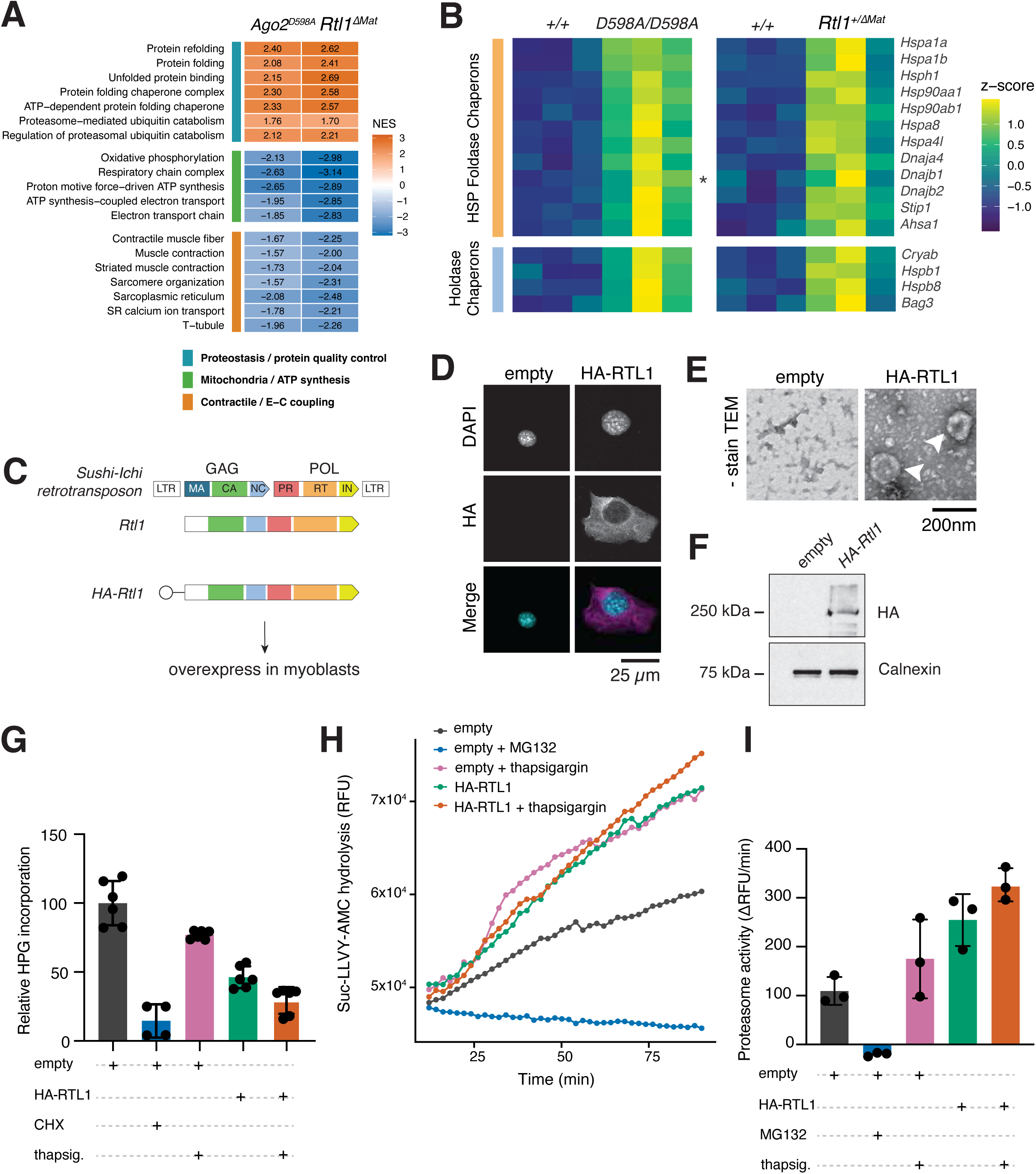
*Rtl1* derepression induces a non-aggregate proteostasis-stress state in skeletal muscle. **(A)** Heatmap of representative GO terms significantly enriched in both *Ago2^D598A/D598A^* and *Rtl1^ΔMat/+^* diaphragm RNA-seq datasets. Values indicate normalized enrichment scores (NES). Positive NES values indicate enrichment among genes upregulated in mutants, whereas negative NES values indicate enrichment among genes downregulated in mutants. **(B)** Heatmap of selected protein-quality-control genes in *Ago2^D598A/D598A^*and *Rtl1^ΔMat/+^* diaphragms. Values represent row z-scores calculated within each dataset. Each column represents an individual biological replicate. All genes are significantly upregulated in both mutants except for Dnajb1 (highlighted with *) which does not reach significance in *Rtl1^ΔMat/+^*diaphragms **(C)** Schematic of the Sushi-ichi retrotransposon, Rtl1, and the HA-Rtl1 expression construct used for overexpression in mouse myoblasts. RTL1 retains retrotransposon-derived Gag- and Pol-related sequences, including a capsid-like CA domain. **(D)** Immunofluorescence of C2C12 myoblasts expressing empty vector or HA-RTL1. **(E)** Negative-stain transmission electron microscopy of virus-like particle (VLP) fractions collected from control or HA-RTL1 expressing cells. **(F)** Immunoblot detection of HA-RTL1 in VLP-enriched fractions isolated from control or HA-RTL1-expressing cells. Calnexin is shown as a control. **(G)** Quantification of nascent protein synthesis by HPG incorporation in C2C12 cells expressing empty vector or HA-RTL1, with cycloheximide (CHX) and thapsigargin treatments as indicated. Values are normalized to empty-vector controls. Each dot represents an independent replicate; bars show mean ± SD. **(H)** Time course of Suc-LLVY-AMC hydrolysis in myoblast cells expressing empty vector or HA-RTL1 and treated as indicated. MG132 serves as a proteasome-inhibition control. RFU, relative fluorescence units. **(I)** Quantification of chymotrypsin-like proteasome activity from the linear phase of the Suc-LLVY-AMC hydrolysis assay in (H). Proteasome activity was quantified as the slope of substrate hydrolysis and is reported as ΔRFU/min. Each dot represents an independent replicate; bars show mean ± SD.

In addition, the shared negatively enriched programs pointed to a coordinated loss of muscle functional capacity. Downregulation of mitochondrial ATP-synthesis programs, including oxidative phosphorylation, respiratory-chain complex assembly, proton-motive-force-driven ATP synthesis, ATP synthesis-coupled electron transport, and electron transport chain, suggests impaired energetic output (**Figure 7A**). These energetic defects coincided with reduced contractile and excitation-contraction coupling programs, including contractile muscle fiber, muscle contraction, striated muscle contraction, sarcomere organization, sarcoplasmic reticulum, SR calcium ion transport, and T-tubule terms (**Figure 7A**). Thus, this signature points to impaired mitochondrial ATP production, defective calcium handling, and reduced contractile capacity in both mouse models. Together, concomitant upregulation of protein-quality-control programs and downregulation of mitochondrial, calcium-handling, and contractile programs suggest that RTL1 derepression places developing muscle in a proteostasis-stress state associated with loss of functional muscle gene programs.

Maintenance of protein homeostasis, or proteostasis, is essential for normal cellular function and organismal health. Proteostasis is maintained by an integrated network that coordinates molecular chaperones, translational control, and proteolytic systems. Chaperones prevent aberrant folding and aggregation while promoting folding and refolding; translational control regulates the influx of newly synthesized polypeptides; proteolytic systems, including the ubiquitin-proteasome and autophagy pathways, remove damaged, excess, misassembled, or terminally misfolded proteins (Hartl, Bracher et al. 2011). Mutations that increase the load of misfolding-prone or assembly-prone proteins can burden protein-quality-control pathways and induce proteostasis stress, which, when sustained, can compromise cellular function (Klaips, Jayaraj et al. 2018).

To test whether RTL1 overexpression could represent one such burden in muscle cells, we overexpressed epitope-tagged RTL1 in mouse myoblasts (**Figure 7C, D**). Previous work showed that mouse RTL1 can form capsid-like particles when overexpressed, similar to other retrotransposon-derived proteins including Arc and PEG10 (Pastuzyn, Day et al. 2018, Segel, Lash et al. 2021). Consistent with this, negative-stain electron microscopy of virus-like particle (VLP)-enriched fractions from HA-RTL1-expressing myoblast cultures revealed large spherical oligomeric structures (**Figure 7E**) and western blotting of these fractions confirmed the presence of HA-RTL1 protein (**Figure 7F**). Thus, in these myoblasts, RTL1 behaves as an oligomerization-prone cytoplasmic protein capable of forming high-molecular-weight particle-like assemblies. One possible outcome of persistent folding or assembly stress caused by the expression of such an oligomerization-prone protein is the formation of insoluble protein inclusions. However, we did not detect cross-β-rich aggresome-like protein aggregates in RTL1-expressing cells, nor did we observe p62-or ubiquitin-positive inclusions in muscle sections from *Ago2* catalytic-mutant embryos. Thus, RTL1 overexpression appears to induce a proteostasis-stress state that is distinct from a classical aggregate myopathy.

In addition to inducing chaperone expression, cells can respond to proteostasis stress by reducing the production of new folding substrates and/or by increasing protein turnover through proteolytic pathways such as proteasome-dependent degradation (Klaips, Jayaraj et al. 2018). We therefore asked whether RTL1 overexpression elicited either of these functional proteostasis responses. First, to quantify nascent protein synthesis, we pulsed cells with L-homopropargylglycine (HPG), a methionine analog that is incorporated into newly synthesized proteins. RTL1 overexpression significantly reduced basal HPG incorporation, indicating that excess RTL1 attenuates nascent protein synthesis even under unstressed conditions (**Figure 7G**). Thapsigargin, an ER Ca2+-ATPase inhibitor that induces ER/proteostasis stress, further reduced HPG incorporation in both empty-vector and RTL1-expressing cells. Two-way ANOVA showed significant main effects of RTL1 expression (p = 1.3 × 10^-4) and thapsigargin treatment (p = 1.8 × 10^-10), with no significant interaction (p = 0.49), indicating that RTL1 and thapsigargin act largely additively to reduce nascent protein synthesis. Planned comparisons confirmed that RTL1 reduced HPG incorporation both at baseline (p = 0.0010) and after thapsigargin treatment (p = 0.0102). (**Figure 7G**). Next, we asked whether RTL1 also altered proteasome-dependent proteolytic activity. To measure proteasome function, we quantified the rate of Suc-LLVY-AMC hydrolysis, a readout of chymotrypsin-like proteasome activity (Morozov, Astakhova et al. 2022). We measured proteasome activity over time (**Figure 7H**) and quantified hydrolysis rate as the slope of substrate cleavage over the linear phase of the assay (**Figure 7I**). We found that RTL1 overexpression increased Suc-LLVY-AMC hydrolysis to levels comparable to thapsigargin-treated controls, which was only modestly increased by additional proteostasis stress (p = 0.12, two-tailed t-test).

Together, our data indicate that unrestrained RTL1 expression induces a functional proteostasis-stress state characterized by basal suppression of nascent protein synthesis and increased proteasome-dependent activity, which can be exacerbated by additional proteostasis challenges. We propose that, in developing diaphragm muscle, where contractile, calcium-handling, mitochondrial, and cytoskeletal proteins must be produced, folded, assembled, and maintained at high demand, this shift in the balance between protein synthesis and protein turnover compromises the maturation and maintenance of the functional muscle proteome, contributing to the lethal muscle failure that characterizes both *Ago2^D598A/D598A^*and *Rtl1^ΔMat/+^*mice.

## DISCUSSION

Befitting Argonaute’s ancestral role in restricting endogenous repeats, we found that Ago2 catalytic-mutant animals have widespread defects caused by failure to repress the domesticated retrotransposon-derived gene *Rtl1*. Our work extends the known impact of AGO2 slicing activity in mammals: in addition to its established role in erythropoiesis through maturation of two noncanonical miRNAs (Cheloufi, Dos Santos et al. 2010, Jee, Yang et al. 2018) AGO2 catalytic activity is required to prevent endothelial and skeletal muscle dysfunction through direct cleavage of the *Rtl1* transcript. The evidence that RTL1 derepression is the major driver of these phenotypes is supported by three lines of evidence. First, *Ago2* catalytic mutants and animals lacking the miRNAs that direct *Rtl1* slicing show overlapping vascular and skeletal muscle defects. Second, our transcriptomic profiling of these two mouse models shows that the gene-level and pathway-level signatures of *Ago2^D598A/D598A^* mutants are largely recapitulated in *Rtl1^ΔMat/+^* animals for both tissues. And third, RTL1 overexpression in myoblasts is sufficient to reproduce key features of the proteostasis-stress response observed in mutant muscle. It is noteworthy that *Rtl1* is an imprinted gene, given that imprinting is thought to have evolved as a mechanism to balance gene dosage (Ishida and Moore 2013). Thus, *Rtl1* expression—like that of other imprinted genes—may need to be kept within a narrow window to remain compatible with mammalian physiology. In the case of *Rtl1*, this dose sensitivity appears to be managed not only through imprinting, but also through direct cleavage of its mRNA by AGO2. In the absence of slicing, *Rtl1* levels become incompatible with normal development of the endothelium and skeletal musculature, with dysfunction of the latter causing postnatal lethality.

In skeletal muscle, the need to tightly regulate *Rtl1* expression may be intimately linked to its evolutionary origin. *Rtl1* is derived from a Ty3/gypsy-type LTR retrotransposon and retains Gag- and Pol-derived sequences, including a capsid-like CA domain (Lynch and Tristem 2003, Youngson, Kocialkowski et al. 2005). Proteins with retrotransposon-derived Gag/CA sequences often retain intrinsic self-assembly capacity, as demonstrated for PEG10, Arc, and mouse RTL1 all of which can assemble into capsid-like particles when overexpressed (Butler, Goodwin et al. 2001, Lynch and Tristem 2003, Segel, Lash et al. 2021). This property may impose an unusual cytoplasmic folding and assembly burden on cells. Consistent with this model, diaphragms from both *Ago2* catalytic mutants and animals lacking the miRNAs that direct *Rtl1* slicing share a heat-shock/protein-quality-control transcriptional response. Moreover, RTL1 overexpression in myoblasts increases MG132-sensitive proteasome activity and reduces nascent protein synthesis, indicating that excess RTL1 is sufficient to functionally perturb proteostasis.

Skeletal muscle may be particularly vulnerable to chronic RTL1-induced proteostasis stress because myofibers must continuously synthesize, fold, assemble, and maintain large sarcomeric, cytoskeletal, mitochondrial, and calcium-handling protein complexes (Hartl, Bracher et al. 2011). In addition, contraction imposes mechanical stress on muscle proteins, requiring chaperone-assisted quality-control pathways, including BAG3/HSPB8/HSPA8-dependent chaperone-assisted selective autophagy, to recognize and clear damaged clients (Ulbricht, Gehlert et al. 2015, Ottensmeyer, Esch et al. 2024). Muscle’s high metabolic and contractile activity also generates reactive oxygen and nitrogen species that, when elevated, can damage proteins and other macromolecules (Powers, Ji et al. 2011). Thus, skeletal muscle is under continuous proteostasis demand even under physiological conditions. In this context, excess RTL1 may impair muscle not by forming classical insoluble aggregates, but by imposing a chronic, non-aggregate cytoplasmic assembly burden that diverts proteostasis capacity away from the maturation and maintenance of the neonatal contractile proteome. This interpretation is in line with the current view that it is the abundance of soluble, aggregate-prone protein oligomers and not that of protein aggregates that drive disease (Henning and Brundel 2017).

Both *Rtl1* and the miRNAs that direct its cleavage are conserved in placental mammals (Edwards, Mungall et al. 2008, Kaneko-Ishino and Ishino 2012, Kozomara and Griffiths-Jones 2014). As a result, we expect that this slicing interaction may have conserved roles in endothelial and skeletal muscle cells across eutherians, representing an additional physiological requirement for slicing that likely contributed to the retention of a catalytically competent Argonaute. In line with this, a companion study showed that overexpression of either mouse *Rtl1* or human RTL1 in C2C12 myoblasts induces overlapping transcriptomic changes, including shared activation of protein-folding/unfolded-protein-response-associated programs (Nickolas Almodovar 2026). Thus, the conserved AGO2–Rtl1 regulatory axis may protect placental mammals from cellular toxicity caused by excess RTL1 dosage.

It is notable that individuals with Kagami-Ogata syndrome (KOS; OMIM #608149)—a disorder caused by genetic or epigenetic disruption of the maternal allele at the 14q32 imprinted locus that contains RTL1—present with many of the features we describe here including RTL1 overexpression, omphalocele, placentomegaly, and respiratory distress (Kagami, Nishimura et al. 2005, Ogata and Kagami 2016, Prasasya, Grotheer et al. 2020) (**Supplementary Table 4**). These parallels suggest that disruption of the conserved RTL1 dosage-control pathway may have important implications for human health. Approximately 25% of individuals diagnosed with KOS die in early childhood, with respiratory complications representing a major cause of mortality; affected individuals can also present with cardiovascular malformations (Kagami, Kurosawa et al. 2015, Ogata and Kagami 2016). Our data raise the possibility that at least some respiratory and cardiovascular manifestations of RTL1 overexpression syndromes arise from RTL1-driven dysfunction in developing muscle and endothelial tissues. In skeletal muscle, this dysfunction may involve a non-aggregate proteostasis-stress state that compromises the maturation and maintenance of the neonatal contractile proteome, suggesting that strategies aimed at preserving proteostasis could be therapeutically relevant.

## ACKNOWLEDGMENTS

We thank all members of the Vidigal lab and of LBMB for discussions and comments on this work. We thank Pedro Rocha, Alex Kelly, Michael Lichten, Sarah Sheppard, Katie McJunkin, Karl Pfeifer, and Todd Macfarlan for critical review of this manuscript. We thank the NCI’s Laboratory Animal Sciences Program, in particular D. Gallardo and M. Figueroa, for expert mouse care and help maintaining the mouse colonies. We also thank the NCI’s molecular histopathology core, especially B. Karim and the Mutant Mouse Resource & Research Center (MMRRC) at UC Davis, especially Louise Lanoue for invaluable pathology work. We also thank Ru-ching Hsia from NCI’s Electron Microscopy Core. This work utilized the computational resources of the NIH HPC Biowulf cluster (<u>hpc.nih.gov</u>). This work was supported by the Intramural Research Program of the National Institutes of Health through the Center for Cancer Research, National Cancer Institute, project 1ZIABC011810 (J.A.V.).

## AUTHOR CONTRIBUTIONS

MK and JAV conceived the project. MK and AGM performed the experiments and analyzed the data with help from MP, AM, JC, and JAV. MK, KF, and HS collected and sequenced *Rtl1^ΔMat/+^* and *Rtl1^+/+^* samples. JAV supervised the project. JAV wrote the manuscript with input from all authors.

## METHODS

### Mouse husbandry and lines

All animal procedures conducted under Animal Study Proposal no. 390923 approved by the Animal Care and Use Committee of the National Cancer Institute or approved by the Animal Ethics Committees of Tokyo Medical and Dental University. Animals were maintained under specific pathogen-free conditions on a 12 h light/12 h dark cycle and had ad libitum access to standard chow and water. *Ago2^D598A^* mice—carrying a point mutation that converts the aspartic acid at position 598 to alanine inactivating AGO2’s catalytic tetrad—(Cheloufi, Dos Santos et al. 2010), *Ago2^flx^*mice—carrying loxP sites around exons 9-11 of the Ago2 gene—(O’Carroll, Mecklenbrauker et al. 2007), Sox2-Cre—which express Cre recombinase under the control of the Sox2 promoter—(Hayashi, Lewis et al. 2002), Tie2-Cre mice—which express Cre recombinase under the control of the Tie2 promoter—(Kisanuki, Hammer et al. 2001) and Acta1-Cre—which express Cre recombinase gene under the control of the human alpha-skeletal actin promoter (Miniou, Tiziano et al. 1999)—were purchased from the Jackson laboratory (Stock nos. 014151, 016520, 008454, 008863, 006149 respectively). The *Rtl1^ΔMat/+^* mice were generated by using ES cells (CCE) derived from the 129/SvEv strain, as previously described (Sekita, Wagatsuma et al. 2008). The *Rtl1^ΔMat/+^*line was maintained by continuous backcrossing with wild-type C57BL/6J mice, and animals from the F10 generation were used in this study. Weaned mice of all strains were genotyped using DNA extracted from tail clippings. Embryos were genotyped using DNA extracted from yolk-sacs.

### µCT analysis

µCT analysis was performed at the Mutant Mouse Resource & Research Center (MMRRC) at UC Davis. A total of 2 *Ago2^+/+^* and 9 *Ago2^D598A/D598A^* littermate animals were processed and analyzed by a trained pathologist. Both genotypes were compared to data from C57BL/6N wild-type mice.

### Immunofluorescence

Tissues were collected and fixed in 10% neutral buffered formalin overnight at room temperature and embedded in paraffin. 5-µm thick tissue sections were dewaxed in xylene and rehydrated through an ethanol series. Sodium citrate buffer [10mM sodium citrate (Millipore Sigma #130-097-418) was used for antigen retrieval, and sections were blocked in blocking buffer [(3% FBS (VWR #97068-091), 1% BSA (Millipore Sigma #A9647), 0.3% Triton X-100 (Millipore Sigma #T8787) in PBS)] prior to incubation with primary antibodies overnight at 4°C. Sections were washed (0.3% Triton X-100 in PBS) and a fluorescence-conjugated secondary antibodies were added for 1 h. After washing, sections were counterstained with DAPI. Sections were mounted with ProLong Gold Antifade reagent. Images were collected using a Axioscan slide scanner (Zeiss Axioscan 7) and analyzed using HALO image analysis platform. Cells were grown on ethanol-sterilized glass coverslips and fixed with 4% paraformaldehyde (PFA) on ice for 10 min. Blocking and permeabilization was carried out for 1 h with PBS containing 5% FCS and 0.3% Triton X-100. The primary antibody was incubated overnight at 4 °C. Coverslips were washed 3x and then incubated for 1 hr in the dark at room temperature with the secondary antibody. All washes occurred for 10 minutes using PBS and all further incubations and washes occurred in the dark. The coverslips were again washed 3x and DNA was counterstained with DAPI, followed by washing 2x. Coverslips were mounted with ProLong Glass Antifade Mountant (Thermo Fisher Scientific). Images were acquired using a Nikon CSU-W1 spinning disk confocal microscope equipped with Photometrics BSI sCMOS camera and ×60 apochromat TIRF (N.A. 1.49) oil immersion lens. Z-stacks were acquired using 0.4 μm *z*-step size and maximum intensity projections were compiled using ImageJ.

### Flow cytometry analysis

*Ago2^+/+^* and *Ago2^D598A/D598A^* placentas were isolated from timed pregnancies of *Ago2^D598A/+^* intercrosses at E16.5. Tissues were resected and briefly washed in cold PBS, before being minced and enzymatically digested [(2.5mg/mL collagenase A (Millipore Sigma #10103586001) 1U/mL Dispase II (Millipore Sigma #4942078001)] in 1X Hank’s Balanced Salt Solution (HBSS). The cell suspension was then filtered through a 70 µm cell strainer (Corning #431751) followed by 40 µm cell strainer (Corning #431750) and pelleted at 400g for 5 minutes at 4°C. Cells were resuspended in 200 µl cold FACS buffer (3% FBS/1%BSA in PBS) and Fc receptors blocked with anti-CD16/32 clone 2.4G2 antibody (BD Pharmingen #553141) for 10 min on ice. Endothelial cells were labelled using the anti-CD31 antibody (BD Bioscience #740879) for 30 min on ice in the dark. Cells were washed twice with FACS buffer and centrifuged at 220g for 5 min at 4°C. Cell pellets were fixed with 4% PFA (Electron Microscopy Sciences #15714) diluted in PBS for 15 min at room temperature. Cells were further washed with FACS buffer and permeabilized with permeabilization solution (0.3% Triton X-100 in FACS buffer) for 10 min at room temperature. Cells were centrifuged at 220g for 5 min at 4°C and resuspended in 200 µl antibody dilution buffer (0.1% Triton X-100 in FACS buffer) and labelled with anti-Ki67 antibody (Invitrogen #25569880) for 40 min on ice. Cells were washed twice in FACS buffer and resuspended in 400 µl FACS buffer. Finally, 50 µl of CountBright Absolute Counting Beads (Thermo Fisher Scientific #C36950) were added into falcon round bottom tubes with cell strainer cap (BD Falcon #352235) directly prior to analysis on a Sony ID7000 FACS machine. Spectral data was processed using the FlowJo software.

### Detection and sequencing of AGO2 cleavage products

Total RNA was extracted from E18.5 placentas using TRIzol (Thermo Fisher Scientific #15596018). Genomic DNA was removed with TURBO DNA-free kit (Thermo Fisher Scientific #AM1907). For each sample, polyadenylated RNA was isolated from 100 μg of total RNA using Dynabeads mRNA purification kit (Thermo Fisher Scientific #610-06) following manufacturer’s instructions. The purified mRNA was used to ligate 5’ RNA adaptor (100 μM) by mixing T4 RNA ligase buffer (NEB # B0216), 10 mM ATP (NEB # P0756), RNaseOUT (Thermo Fisher Scientific #10777-019), and T4 RNA ligase (NEB # M0204) as previously described (Li, Zhao et al. 2019). A second round of mRNA purification was performed to remove unincorporated 5’ RNA adaptors. Adaptor-ligated mRNA was reverse-transcribed using SuperScript™ II (Thermo Fisher, Cat. No. 18064) by combining transcript-specific primers and a β-actin-specific primer. cDNA was amplified with a forward primer designed to bind to the 5’ RNA adaptor sequence and a gene-specific reverse primer using the high-fidelity Platinum Taq DNA Polymerase (Thermo Fisher Scientific #11304011) supplemented with DMSO (Thermo Fisher Scientific #F-515) and betaine (Thermo Fisher Scientific #J77507.VCR). The PCR cycling conditions were as follows: 94°C for 2 min, 35 cycles of 94°C for 20 s, 60°C for 30 s, and 68°C for 45 s; and 72°C for 7 min.

PCR products were run on 2% agarose gel and extracted using Zymoclean Gel DNA recovery kit (Zymo #D4008) following manufacturer’s instruction. Extracted amplicons were cloned using Zero Blunt TOPO PCR cloning kit (Thermo Fisher Scientific #450245), and plasmid DNA isolated from individual colonies sent for sanger sequencing.

### Quantitative RT-PCR

RNA was isolated using TRIzol reagent (Thermo Fisher Scientific #15596018) and DNase treated with TURBO DNA-free kit (Thermo Fisher Scientific #AM1907) following manufacturers’ protocols. RNA was reverse transcribed with ABScript III RT Master Mix (Abclonal # RK20428) using random hexamers. Relative expression level of *Rtl1* was quantified using Sybr Green (Thermo Fisher Scientific #4309155) and normalized to either *Hprt* or *Gapdh* housekeeping gene. Primer sequences are as follows: *Rtl1* (AGGTTGCAAGCTTGCTGGT, TGAAGTACTCATTATAGTCAAGGGCA), *Hprt* (TGGAGCCACCGATCCACACA, CGTTGACATCCGTAAAGACC), *Gapdh* (CGTCCCGTAGACAAAATGGT, TCAATGAAGGGGTCGTTGAT).

### Western blot

Tissues were homogenized on ice in 500 µl SDS-lysis buffer (2% SDS, 50mM Tris, 10% Glycerol) containing protease inhibitors (Millipore Sigma #PPC2020) and protein concentration was determined using the Pierce BCA protein assay kit (Thermo Scientific #23225). Samples were stored at −80^0^ C until analysis. Protein extracts were mixed with Laemmli sample buffer (Bio-Rad #1610747), boiled for 5 minutes, and chilled briefly on ice prior to loading on gels. Proteins were resolved by NuPAGE™ Bis-Tris Mini Protein Gels, 4–12% (Thermo Fisher Scientific #NP0322BOX) electrophoresis and transferred to nitrocellulose membrane (Thermo Fisher Scientific #88018) by electroblotting. Membranes were blocked in 5% BSA (Millipore Sigma #A9647) in Tris-buffered saline (10 mM Tris, pH 7·5, 150 mM sodium chloride) containing 0.1% Tween 20 (TBST). The p-Histone H3 (Santacruz #sc-8656; dilution 1:500), Cleaved-caspase-3 (Cell Signaling #9661; dilution 1:500), HA-Tag (Cell Signaling #3724; dilution 1:1000), Calnexin (Abcam #ab22595; dilution 1:1000), Histone H3 (Abcam #ab1791; dilution 1:2000), and Gapdh (Santacruz #sc-32233; dilution 1:4000) antibodies were used in TBST, 5% BSA overnight at 4°C. Membranes were washed and anti-Rabbit IgG HRP (Cell Signaling #7074) or anti-mouse IgG HRP (Cell Signaling #7076) were added for 1 h. Proteins were developed using Supersignal West Dura Extended duration substrate (Thermo Fisher Scientific #34076) and visualized by chemiluminescence (Amersham ImageQuant 800 systems).

### Skeletal Preparations

Embryos (E18.5) were eviscerated and soaked in ddH_2_O for 3 h at room temperature (17-22°C) as described previously (de Pontual, Yao et al. 2011). Fetuses were heat shocked in a 65°C water bath for 30 s to facilitate removal of skin. Embryos were fixed in absolute ethanol (Millipore Sigma #E7023) followed by acetone (Millipore Sigma #179124) overnight at room temperature. Cartilages were soaked in alcian blue (Millipore Sigma #A5268) solution (150 mg/l alcian blue 8GX, 80% ethanol, 20% acetic acid) and alizarin red (Millipore Sigma #A5533) solution (50 mg/l alizarin red S in 2% KOH) to stain for cartilage and bone respectively. Remaining tissues were cleared in 2% KOH (Millipore Sigma #105033) dissolved in ddH_2_O, and skeletons were placed in 25% glycerol (Millipore Sigma #G5516) for storage. Images were captured with a Zeiss Stereo Discovery V8 microscope and processed in Photoshop. Measurements of bone length were performed in ImageJ.

### RNA sequencing

For total RNA sequencing (*Ago2* mutant and wild-type placentas; *Rtl1* maternal deletion and wild-type tissues) ribosomal RNA was depleted from extracted total RNA using the Ribo-Zero Plus kit (cat# 20040525) or ZapR® Depletion Kit (Takara Bio, #634360). Libraries were constructed either using the Illumina TruSeq Stranded Total RNA Library (Ago2 samples) or SMART-Seq® Total RNA Single Cell Library Prep (Rtl1 samples) and subjected to paired end sequencing on NextSeq 2000 or NovaSeq X Plus instruments. Reads were trimmed using cutadapt (v 4.7) and mapped to GRCm39 genome using STAR aligner (2.7.11b). Read alignments were counted using Subread featureCounts (v 2.0.3). Differentially expressed genes were identified with DEseq2 (R 3.6.0).

### Small RNA sequencing

Libraries were constructed using NEBNext^®^ Multiplex Small RNA Library Prep Set for Illumina^®^ (cat #E7300S) following manufacturer’s instructions. Sequencing was performed on NextSeq 500 using SR75. Reads were trimmed using cutadapt (v 4.7). Trimmed reads were mapped to either miRNA or hairpin customized genomes using STAR aligner, with sequences retrieved from miRBase (2.7.11b). Read alignments were counted using Subread featureCounts (v 2.0.3) where features were retrieved from miRBase. Differentially expressed mature and precursor miRNAs were identified using DEseq2 (R 3.6.0).

### Cell culture

C2C12 myoblasts were purchased from ATCC and maintained on DMEM media supplemented with 10% FBS. Stable C2C12 cell lines overexpressing HA-tagged RTL1 were generated using the PiggyBac transposon system. C2C12 myoblasts were co-transfected with a PiggyBac transposon vector encoding HA-tagged mouse RTL1 (PB-mRtl1-PGKpuro) and the hyperactive PiggyBac transposase expression plasmid pCMV-hyPBase (pVL283) at a 9:1 DNA ratio, respectively as previously described (Riordan, Keng et al. 2013) using the Lonza Nucleofector system according to the manufacturer’s instructions. Forty-eight hours after nucleofection, cells were subjected to puromycin selection (1 μg/mL). Selection was continued until non-transfected control cells were eliminated and resistant colonies were established.

### Virus-Like Particle (VLP) Isolation

C2C12 cells were seeded in 150 mm dishes in DMEM overnight to reach about 60% confluency. Cells were washed twice with PBS and maintained in conditioned medium, Opti-MEM (Thermo Fisher Scientific, Cat. No. 11058021) for 48 hours prior to VLPs collection. Conditioned medium was clarified by centrifugation at 2,000 × g for 10 min at 4°C and filtered through a 0.45 μm Durapore PVDF filter (EMD Millipore, Cat. No. SE1M003M00). Supernatants were centrifuged at 120,000 × g for 2 h at 4°C using an Optima XL-100K ultracentrifuge (Beckman Coulter) equipped with a Type 45 Ti rotor. Following centrifugation, supernatants were discarded and pellets from all tubes corresponding to the same experimental condition were pooled and resuspended in a final volume of 100 μL PBS for downstream analyses. Transmission electron microscopy analysis of these fractions was performed at NCI’s Electron Microscopy Core.

### Proteasome activity

C2C12 myoblasts were maintained under standard culture conditions and seeded at 7.5 × 10^4 cells per well in 6-well plates. Cells were cultured for 24 h prior to experimental treatments. Where indicated, cells were treated with 1 μM thapsigargin (MedChemExpress, Cat. No. HY-13433) for 4 h. Cells were harvested at approximately 70–90% confluence and processed for proteasome activity assays. Proteasome chymotrypsin-like activity was measured using the fluorogenic substrate Suc-Leu-Leu-Val-Tyr-7-amido-4-methylcoumarin (Suc-LLVY-AMC; AAT Bioquest, Cat. No. 13453). C2C12 cells were washed twice with PBS and detached using ESGRO Complete Accutase Cell Dissociation Reagent (MilliporeSigma, Cat. No. SF006) according to the manufacturer’s instructions. Cells were collected by centrifugation (500 × g, 5 min, 4°C), washed once with ice-cold PBS, immediately transferred to ice, and resuspended in assay buffer containing 10 mM Tris-HCl (pH 7.5), 25 mM KCl, 1.1 mM MgCl2, 0.1 mM EDTA, and 10% glycerol. 1 mM DTT was added to the buffer immediately before use. Cell lysates were generated by three freeze–thaw cycles using a dry ice-ethanol slurry followed by rapid thawing at 37°C. Lysates were clarified by centrifugation at 16,000 × g for 15 min at 4°C. Protein concentrations were determined using a bicinchoninic acid (BCA) assay. Proteasome activity assays were performed in triplicate in black-walled, clear-bottom 96-well microplates (Costar; Corning, Cat. No.3340) in a final reaction volume of 100 μL. Reaction mixtures contained 20 μg whole-cell lysate per well, assay buffer, 0.02% SDS, and Suc-LLVY-AMC substrate at a final concentration of 50 μM. Assay specificity was confirmed using the proteasome inhibitor MG132 (MilliporeSigma, Cat. No. 474791-1MG; 10 μM final concentration). Blank wells lacking lysate were included for background correction. Fluorescence was monitored using a Synergy Neo2 multimode plate reader (BioTek Instruments, Winooski, VT, USA) equipped for top-read fluorescence measurements. Plates were equilibrated at 37°C and subjected to linear shaking for 5 s prior to data acquisition. Kinetic fluorescence measurements were collected for 90 min at 2-min intervals using excitation and emission filters of 400/30 nm and 460/40 nm, respectively. Proteasome activity was quantified from the slope of the linear phase of AMC fluorescence accumulation and is reported as ΔRFU/min.

### Measurement of Nascent Protein Synthesis by HPG Incorporation and Flow Cytometry

Nascent protein synthesis was quantified using the Click-iT™ HPG Alexa Fluor™ 488 Protein Synthesis Assay Kit (Thermo Fisher Scientific, Cat. No. C10428) according to the manufacturer’s instructions with minor modifications for flow cytometric analysis. C2C12 myoblasts were seeded at 3.5 × 10^4 cells per well in 12-well plates and cultured overnight. The following day, cells were either left untreated, treated with 1 μM thapsigargin (MedChemExpress, Cat. No. HY-13433) for 4 h, or treated with cycloheximide (50 μg/mL) for 30 min as a control for protein synthesis inhibition. During the final 30 min of treatment, cells were incubated with L-homopropargylglycine (HPG) in methionine-free DMEM (high glucose, no glutamine, no methionine, and no cystine; Thermo Fisher Scientific, Cat. No. 21013024) according to the manufacturer’s recommendations. Cycloheximide-treated cells remained exposed to cycloheximide throughout the HPG-labeling period. Cells were then detached using ESGRO Complete Accutase Cell Dissociation Reagent (MilliporeSigma, Cat. No. SF006), washed with PBS containing 3% BSA, fixed, and permeabilized according to the Click-iT HPG assay protocol. Incorporated HPG was detected by copper-catalyzed azide–alkyne cycloaddition using the Alexa Fluor 488 azide detection reagent supplied in the kit. Samples were analyzed on a BD FACSymphony flow cytometer (BD Biosciences). Debris and doublets were excluded based on forward- and side-scatter characteristics, and Alexa Fluor 488 fluorescence intensity within the singlet population was used as a measure of nascent protein synthesis. HPG incorporation was normalized to the untreated control group and is presented as relative HPG incorporation.

## Data availability

FASTQ files and processed RNA-seq datasets generated in this study have been deposited in the Gene Expression Omnibus under accession numbers GSE291207, GSE291208, and GSE329372.

---

**Sup. Fig. 1.**
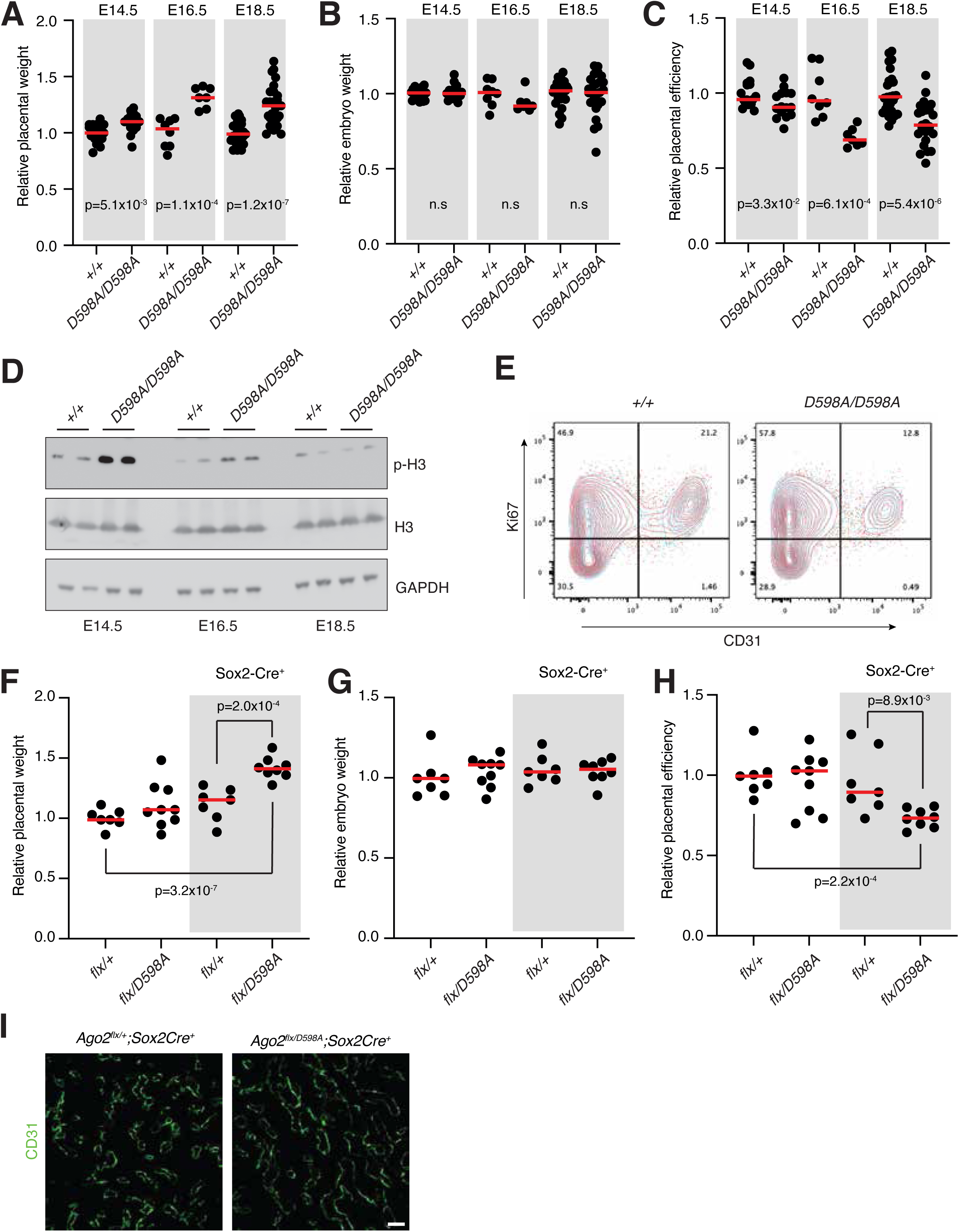
Placentomegaly in *Ago2* catalytic mutants. **(A)** Relative placental weights of mutants relative to wild-type littermates at E18.5. **(B)** Relative embryo weights of mutants relative to wild-type littermates at E18.5. **(C)** Relative placental efficiency of mutants relative to wild-type littermates at E18.5. Each dot represents an animal. p-values were calculated using a two-tailed t-test. **(D)** Histone H3 phosphorylation as measured by western blot on protein lysates from whole E14.5, E16.5, and E18.5 placentas. **(E)** Flow cytometry analysis of E16.5 placentas using antibodies against the proliferative marker Ki67 and the endothelial cell marker CD31. Plots show overlay of two biological replicates per genotype, each highlighted with a different color. **(F**, **G**, **H)** As (A, B, C) but for embryos from the Sox2-Cre crosses. **(I)** immunofluorescence staining of mouse placentas with an antibody against the CD31 endothelial cell marker. For all graphs each dot represents an animal. p-values were calculated using a two-tailed t-test.

**Sup. Fig. 2.**
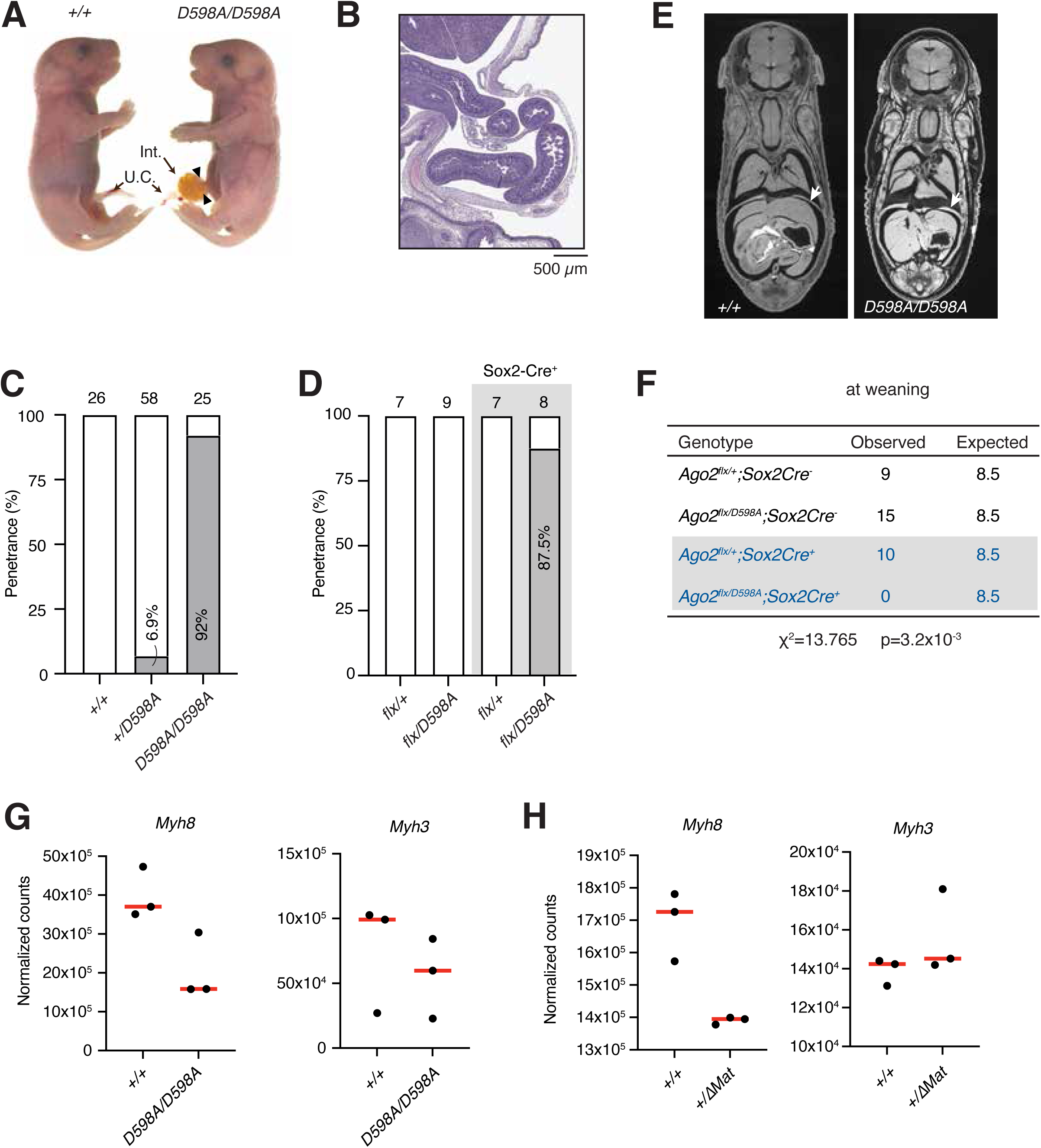
Characterization of Ago2 slicing mutants. **(A)** Gross appearance of wild-type (+/+) and catalytic mutant (D598A/D598A) animals at E18.5. Int., intestine; U.C., umbilical cord. **(B)** Hematoxylin-Eosin-stained sagittal section of omphalocele. **(C)** Penetrance of omphalocele in E18.5 animals. **(D)** Penetrance of omphalocele in E18.5 animals from Sox2-Cre crosses. **(E)** Micro-CT frontal sections of wild-type (+/+) and catalytic mutant (D598A/D598A) animals at E18.5. Note the thickened diaphragm muscle (arrow). **(F)** Genotyping of litters at weaning showing conditional loss of AGO2 catalytic competence in the Sox2+ cell lineage is results postnatal lethality. P value was calculated with the Chi-square test. **(G, H)** Normalized counts for indicated genes in *Ago2* (G) and *Rtl1* (H) sequencing datasets.

**Sup. Fig. 3.**
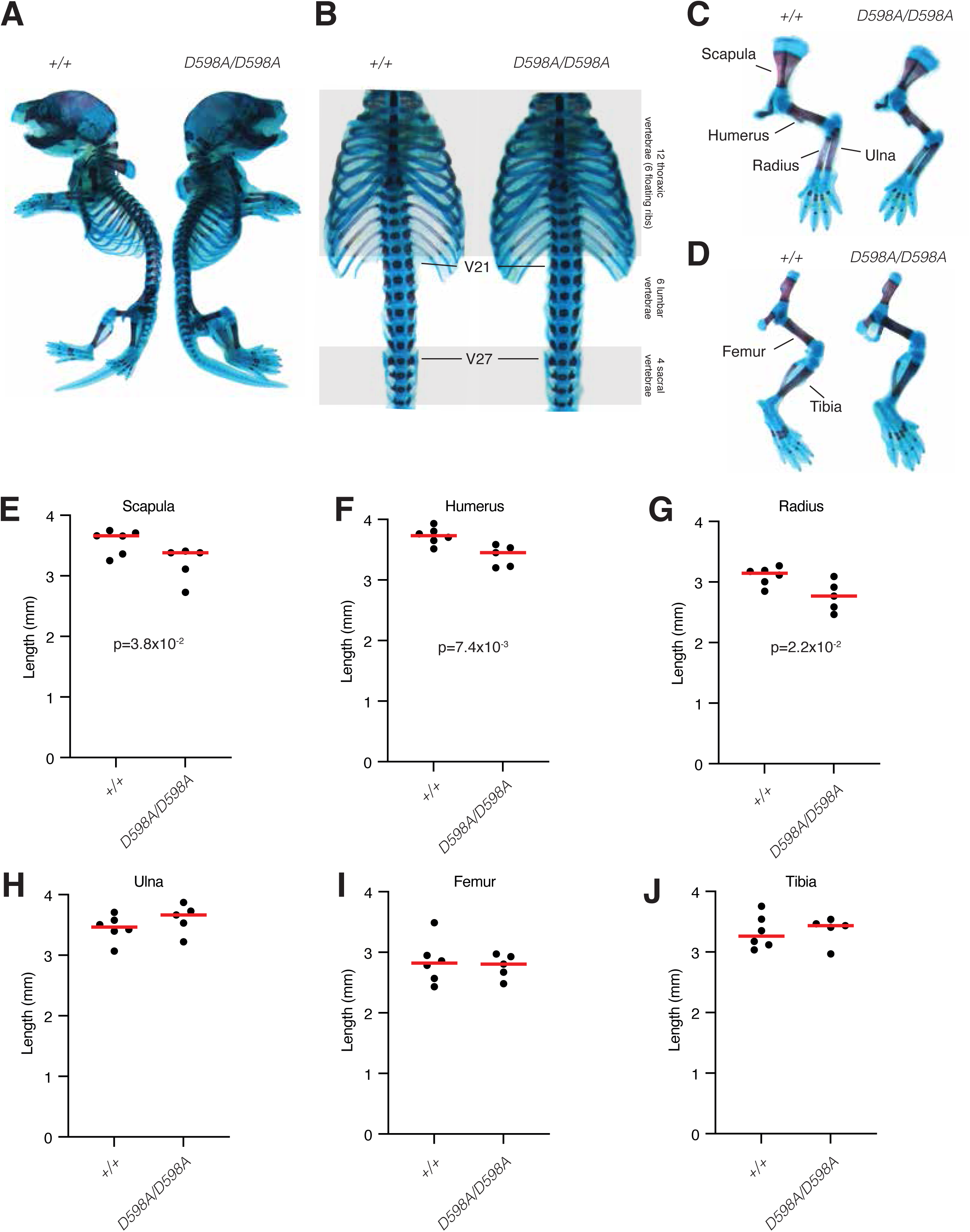
Skeletal development in the absence of AGO2 catalytic activity. **(A-D)** Representative examples of skeletons from E18.5 wild-type and homozygous mutant animals stained with alcian blue (for cartilage) and alizarin red (for bone). The positions of vertebra 21 (V21) and 27 (V27) which correspond to the first lumbar and first sacral vertebras in both genotypes are highlighted in (B). **(E-J)** Length of the bones highlighted in (C-D) for each genotype. Each dot represents an animal. P-values were calculated using a two-tailed t-test.

**Sup. Fig. 4.**
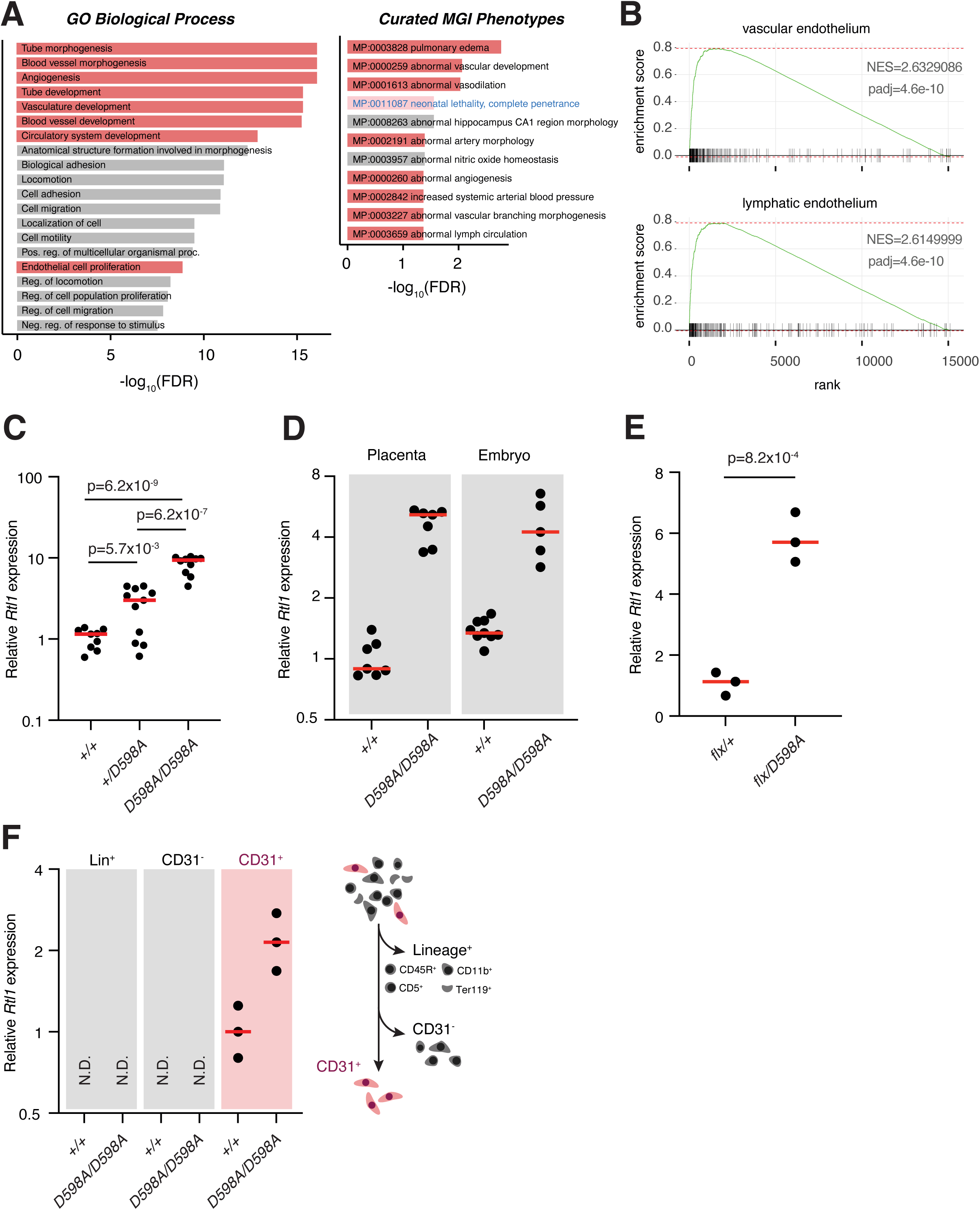
Endothelial cell dysregulation in AGO2 catalytic mutant mice. **(A)** Enriched GO terms (left) and MGI Phenotypes (right) among genes upregulated in Ago2 catalytic mutant placentas. Terms related to vascular development are highlighted in red. Neonatal lethality is highlighted in pink. **(B)** Gene Set Enrichment Analysis (GSEA) plots showing a significant enrichment of gene signatures for vascular and lymphatic endothelium amongst the genes dysregulated in *Ago2* catalytic mutants. (C) Relative *Rtl1* expression in E18.5 placentas of the indicated genotypes **(D)** Relative *Rtl1* expression in E14.5 placentas and embryos. All values are normalized to the average of the wild-type placenta. Each dot represents a biological replicate. P-values were calculated using a two-tailed t-test. (**E)** Relative *Rtl1* expression from *Ago2^flx/+^;Sox2-Cre^+^* and *Ago2^flx/D598A^;Sox2-Cre^+^* E18.5 placentas. Each dot represents a biological replicate. P-values were calculated using a two-tailed t-test. **(F)** Relative *Rtl1* expression in CD31+ endothelial cells isolated from E14.5 embryos. Each dot represents an animal. Schematic representation of the isolated fractions is shown on the right.

**Sup. Fig. 5.**
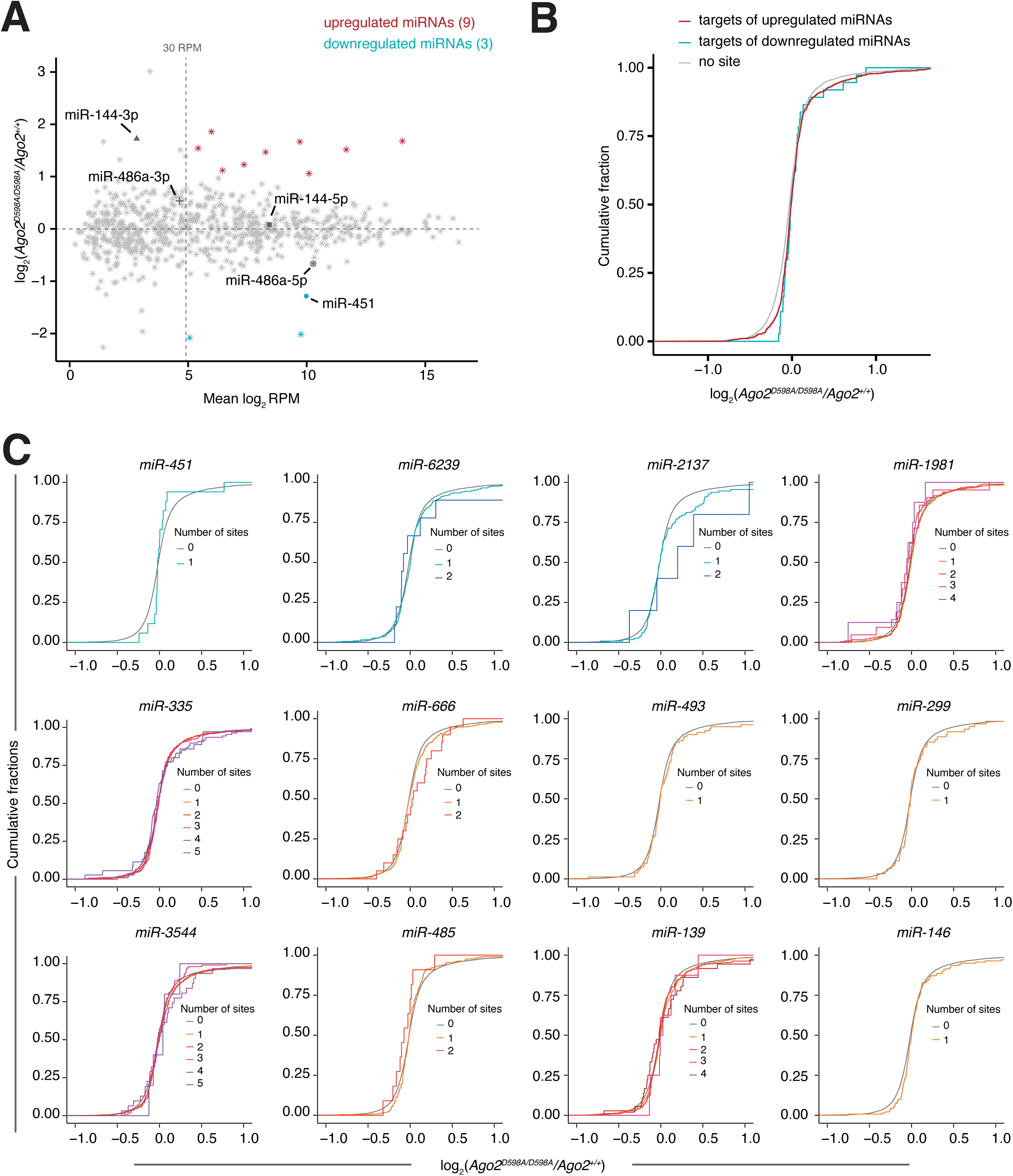
Transcriptomic changes are not caused by miRNA dysregulation. **(A)** Analysis of small RNA libraries in wild-type and mutant placentas. We consider well-expressed miRNAs to be those with more than 30 RPM in these datasets. The biogenesis of *miR-486* and *miR-451* was previously shown to depend on AGO2 slicing activity, and both are highlighted in the plot along with *miR-144*, which is co-transcribed with *miR-451*. **(B)** Cumulative fraction plots showing that the expression of predicted targets of dysregulated miRNAs is not preferentially impacted. **(C)** As in (B) but highlighting the targets of individual downregulated (blue tones) or upregulated (red tones) miRNAs by number of predicted target sites.

**Sup. Fig. 6.**
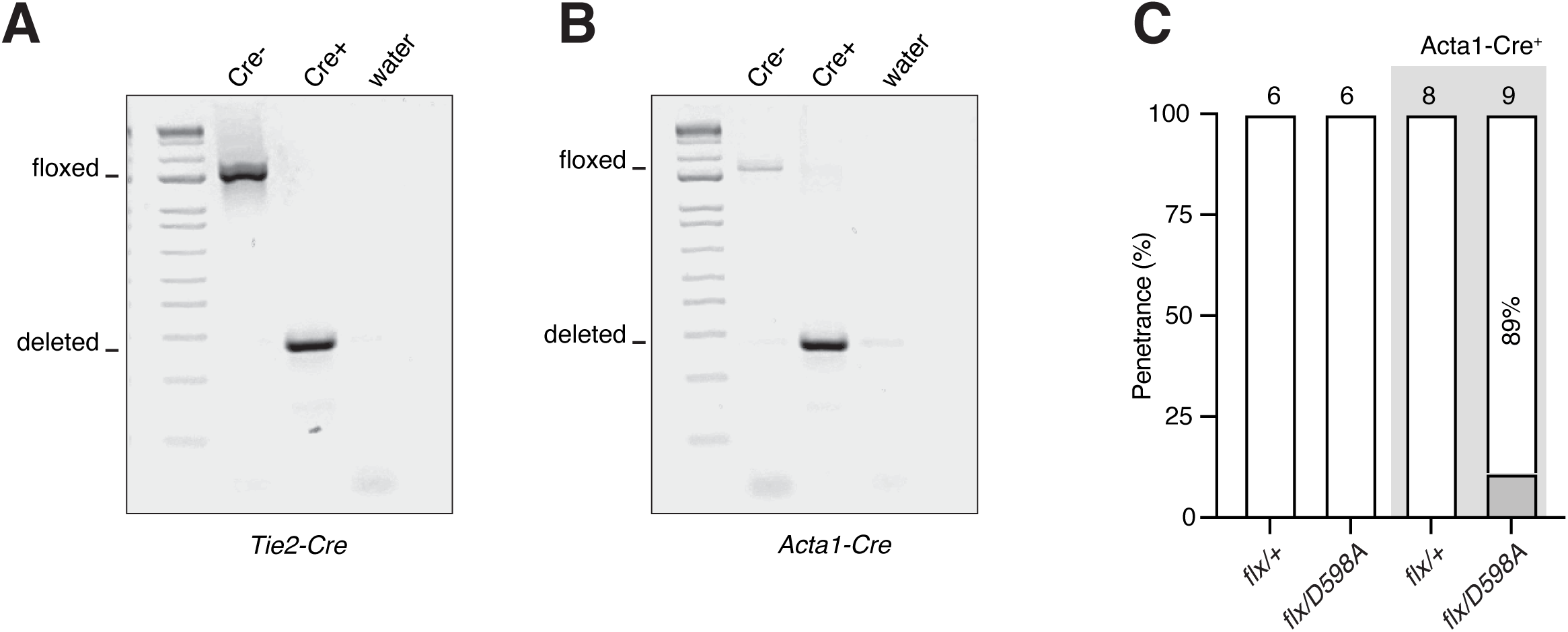
Conditional deletion of the Ago2 floxed allele in endothelial cells and skeletal muscle. **(A**, **B)** Genotyping PCR showing deletion of the floxed allele in CD31+ cells of *Tie2-Cre+* embryos (A) and in the skeletal muscle of *Acta1-Cre+* embryos (B). **(C)** Penetrance of omphalocele in E18.5 animals from the *Acta1-Cre* crosses.

## REFERENCES

Bartel, D. P. (2018). “Metazoan MicroRNAs.” Cell 173(1): 20–51.

Becker, W. R., B. Ober-Reynolds, K. Jouravleva, S. M. Jolly, P. D. Zamore and W. J. Greenleaf (2019). “High-Throughput Analysis Reveals Rules for Target RNA Binding and Cleavage by AGO2.” Mol Cell 75(4): 741–755 e711.

Bourque, G., K. H. Burns, M. Gehring, V. Gorbunova, A. Seluanov, M. Hammell, M. Imbeault, Z. Izsvak, H. L. Levin, T. S. Macfarlan, D. L. Mager and C. Feschotte (2018). “Ten things you should know about transposable elements.” Genome Biol 19(1): 199.

Burns, K. H. (2022). “Repetitive DNA in disease.” Science 376(6591): 353–354.

Butler, M., T. Goodwin, M. Simpson, M. Singh and R. Poulter (2001). “Vertebrate LTR retrotransposons of the Tf1/sushi group.” J Mol Evol 52(3): 260–274.

Cavaille, J., H. Seitz, M. Paulsen, A. C. Ferguson-Smith and J. P. Bachellerie (2002). “Identification of tandemly-repeated C/D snoRNA genes at the imprinted human 14q32 domain reminiscent of those at the Prader-Willi/Angelman syndrome region.” Hum Mol Genet 11(13): 1527–1538.

Chapman, E., F. Taglini and E. H. Bayne (2022). “Separable roles for RNAi in regulation of transposable elements and viability in the fission yeast Schizosaccharomyces japonicus.” PLoS Genet 18(2): e1010100.

Charite, J., W. de Graaff, S. Shen and J. Deschamps (1994). “Ectopic expression of Hoxb-8 causes duplication of the ZPA in the forelimb and homeotic transformation of axial structures.” Cell 78(4): 589–601.

Charlier, C., K. Segers, D. Wagenaar, L. Karim, S. Berghmans, O. Jaillon, T. Shay, J. Weissenbach, N. Cockett, G. Gyapay and M. Georges (2001). “Human-ovine comparative sequencing of a 250-kb imprinted domain encompassing the callipyge (clpg) locus and identification of six imprinted transcripts: DLK1, DAT, GTL2, PEG11, antiPEG11, and MEG8.” Genome Res 11(5): 850–862.

Cheloufi, S., C. O. Dos Santos, M. M. Chong and G. J. Hannon (2010). “A dicer-independent miRNA biogenesis pathway that requires Ago catalysis.” Nature 465(7298): 584–589.

Chung, W. J., K. Okamura, R. Martin and E. C. Lai (2008). “Endogenous RNA interference provides a somatic defense against Drosophila transposons.” Curr Biol 18(11): 795–802.

Davis, E., F. Caiment, X. Tordoir, J. Cavaille, A. Ferguson-Smith, N. Cockett, M. Georges and C. Charlier (2005). “RNAi-mediated allelic trans-interaction at the imprinted Rtl1/Peg11 locus.” Curr Biol 15(8): 743–749.

de Pontual, L., E. Yao, P. Callier, L. Faivre, V. Drouin, S. Cariou, A. Van Haeringen, D. Genevieve, A. Goldenberg, M. Oufadem, S. Manouvrier, A. Munnich, J. A. Vidigal, M. Vekemans, S. Lyonnet, A. Henrion-Caude, A. Ventura and J. Amiel (2011). “Germline deletion of the miR-17 approximately 92 cluster causes skeletal and growth defects in humans.” Nat Genet 43(10): 1026–1030.

Edwards, C. A., A. J. Mungall, L. Matthews, E. Ryder, D. J. Gray, A. J. Pask, G. Shaw, J. A. Graves, J. Rogers, S. consortium, I. Dunham, M. B. Renfree and A. C. Ferguson-Smith (2008). “The evolution of the DLK1-DIO3 imprinted domain in mammals.” PLoS Biol 6(6): e135.

Fire, A., S. Xu, M. K. Montgomery, S. A. Kostas, S. E. Driver and C. C. Mello (1998). “Potent and specific genetic interference by double-stranded RNA in Caenorhabditis elegans.” Nature 391(6669): 806–811.

Hartl, F. U., A. Bracher and M. Hayer-Hartl (2011). “Molecular chaperones in protein folding and proteostasis.” Nature 475(7356): 324–332.

Hayashi, S., P. Lewis, L. Pevny and A. P. McMahon (2002). “Efficient gene modulation in mouse epiblast using a Sox2Cre transgenic mouse strain.” Mech Dev 119 **Suppl 1**: S97–S101.

He, X., Y. L. Yan, J. K. Eberhart, A. Herpin, T. U. Wagner, M. Schartl and J. H. Postlethwait (2011). “miR-196 regulates axial patterning and pectoral appendage initiation.” Dev Biol 357(2): 463–477.

Henning, R. H. and B. Brundel (2017). “Proteostasis in cardiac health and disease.” Nat Rev Cardiol 14(11): 637–653.

Ishida, M. and G. E. Moore (2013). “The role of imprinted genes in humans.” Mol Aspects Med 34(4): 826–840.

Jee, D., J. S. Yang, S. M. Park, D. T. Farmer, J. Wen, T. Chou, A. Chow, M. T. McManus, M. G. Kharas and E. C. Lai (2018). “Dual Strategies for Argonaute2-Mediated Biogenesis of Erythroid miRNAs Underlie Conserved Requirements for Slicing in Mammals.” Mol Cell 69(2): 265–278 e266.

Jonas, S. and E. Izaurralde (2015). “Towards a molecular understanding of microRNA-mediated gene silencing.” Nat Rev Genet 16(7): 421–433.

Joureau, B., J. M. de Winter, K. Stam, H. Granzier and C. A. Ottenheijm (2017). “Muscle weakness in respiratory and peripheral skeletal muscles in a mouse model for nebulin-based nemaline myopathy.” Neuromuscul Disord 27(1): 83–89.

Kagami, M., K. Kurosawa, O. Miyazaki, F. Ishino, K. Matsuoka and T. Ogata (2015). “Comprehensive clinical studies in 34 patients with molecularly defined UPD(14)pat and related conditions (Kagami-Ogata syndrome).” Eur J Hum Genet 23(11): 1488–1498.

Kagami, M., G. Nishimura, T. Okuyama, M. Hayashidani, T. Takeuchi, S. Tanaka, F. Ishino, K. Kurosawa and T. Ogata (2005). “Segmental and full paternal isodisomy for chromosome 14 in three patients: narrowing the critical region and implication for the clinical features.” Am J Med Genet A 138A(2): 127–132.

Kaneko-Ishino, T. and F. Ishino (2012). “The role of genes domesticated from LTR retrotransposons and retroviruses in mammals.” Front Microbiol 3: 262.

Kazazian, H. H., Jr. and J. V. Moran (2017). “Mobile DNA in Health and Disease.” N Engl J Med 377(4): 361–370.

Kisanuki, Y. Y., R. E. Hammer, J. Miyazaki, S. C. Williams, J. A. Richardson and M. Yanagisawa (2001). “Tie2-Cre transgenic mice: a new model for endothelial cell-lineage analysis in vivo.” Dev Biol 230(2): 230–242.

Kitazawa, M., S. Hayashi, M. Imamura, S. Takeda, Y. Oishi, T. Kaneko-Ishino and F. Ishino (2020). “Deficiency and overexpression of Rtl1 in the mouse cause distinct muscle abnormalities related to Temple and Kagami-Ogata syndromes.” Development 147(21).

Klaips, C. L., G. G. Jayaraj and F. U. Hartl (2018). “Pathways of cellular proteostasis in aging and disease.” J Cell Biol 217(1): 51–63.

Klattenhoff, C., D. P. Bratu, N. McGinnis-Schultz, B. S. Koppetsch, H. A. Cook and W. E. Theurkauf (2007). “Drosophila rasiRNA pathway mutations disrupt embryonic axis specification through activation of an ATR/Chk2 DNA damage response.” Dev Cell 12(1): 45–55.

Kozomara, A. and S. Griffiths-Jones (2014). “miRBase: annotating high confidence microRNAs using deep sequencing data.” Nucleic Acids Res 42(Database issue): D68–73.

Li, Y. F., M. Zhao, M. Wang, J. Guo, L. Wang, J. Ji, Z. Qiu, Y. Zheng and R. Sunkar (2019). “An improved method of constructing degradome library suitable for sequencing using Illumina platform.” Plant Methods 15: 134.

Liu, J., M. A. Carmell, F. V. Rivas, C. G. Marsden, J. M. Thomson, J. J. Song, S. M. Hammond, L. Joshua-Tor and G. J. Hannon (2004). “Argonaute2 is the catalytic engine of mammalian RNAi.” Science 305(5689): 1437–1441.

Lynch, C. and M. Tristem (2003). “A co-opted gypsy-type LTR-retrotransposon is conserved in the genomes of humans, sheep, mice, and rats.” Curr Biol 13(17): 1518–1523.

Maksakova, I. A., D. L. Mager and D. Reiss (2008). “Keeping active endogenous retroviral-like elements in check: the epigenetic perspective.” Cell Mol Life Sci 65(21): 3329–3347.

Marsh, B. and R. Blelloch (2020). “Single nuclei RNA-seq of mouse placental labyrinth development.” Elife 9.

Matzke, M. A. and J. A. Birchler (2005). “RNAi-mediated pathways in the nucleus.” Nat Rev Genet 6(1): 24–35.

Miniou, P., D. Tiziano, T. Frugier, N. Roblot, M. Le Meur and J. Melki (1999). “Gene targeting restricted to mouse striated muscle lineage.” Nucleic Acids Res 27(19): e27.

Morozov, A., T. Astakhova, P. Erokhov and V. Karpov (2022). “The ATP/Mg(2+) Balance Affects the Degradation of Short Fluorogenic Substrates by the 20S Proteasome.” Methods Protoc 5(1).

Nakanishi, K., D. E. Weinberg, D. P. Bartel and D. J. Patel (2012). “Structure of yeast Argonaute with guide RNA.” Nature 486(7403): 368–374.

Nichol, P. F., R. F. Corliss, J. D. Tyrrell, B. Graham, A. Reeder and Y. Saijoh (2011). “Conditional mutation of fibroblast growth factor receptors 1 and 2 results in an omphalocele in mice associated with disruptions in ventral body wall muscle formation.” J Pediatr Surg 46(1): 90–96.

Nickolas Almodovar, C. D., Benjamin Kleaveland (2026). “Argonaute-2 slicing of Rtl1 promotes skeletal muscle development and postnatal survival.” bioRxiv.

O’Carroll, D., I. Mecklenbrauker, P. P. Das, A. Santana, U. Koenig, A. J. Enright, E. A. Miska and A. Tarakhovsky (2007). “A Slicer-independent role for Argonaute 2 in hematopoiesis and the microRNA pathway.” Genes Dev 21(16): 1999–2004.

Obbard, D. J., K. H. Gordon, A. H. Buck and F. M. Jiggins (2009). “The evolution of RNAi as a defence against viruses and transposable elements.” Philos Trans R Soc Lond B Biol Sci 364(1513): 99–115.

Ogata, T. and M. Kagami (2016). “Kagami-Ogata syndrome: a clinically recognizable upd(14)pat and related disorder affecting the chromosome 14q32.2 imprinted region.” J Hum Genet 61(2): 87–94.

Ottensmeyer, J., A. Esch, H. Baeta, S. Sieger, Y. Gupta, M. F. Rathmann, A. Jeschke, D. Jacko, K. Schaaf, T. Schiffer, B. Rahimi, L. Lovenich, A. Sisto, P. F. M. van der Ven, D. O. Furst, A. Haas, W. Bloch, S. Gehlert, B. Hoffmann, V. Timmerman, P. F. Huesgen and J. Hohfeld (2024). “Force-induced dephosphorylation activates the cochaperone BAG3 to coordinate protein homeostasis and membrane traffic.” Curr Biol 34(18): 4170–4183 e4179.

Pastuzyn, E. D., C. E. Day, R. B. Kearns, M. Kyrke-Smith, A. V. Taibi, J. McCormick, N. Yoder, D. M. Belnap, S. Erlendsson, D. R. Morado, J. A. G. Briggs, C. Feschotte and J. D. Shepherd (2018). “The Neuronal Gene Arc Encodes a Repurposed Retrotransposon Gag Protein that Mediates Intercellular RNA Transfer.” Cell 173(1): 275.

Perez-Garcia, V., E. Fineberg, R. Wilson, A. Murray, C. I. Mazzeo, C. Tudor, A. Sienerth, J. K. White, E. Tuck, E. J. Ryder, D. Gleeson, E. Siragher, H. Wardle-Jones, N. Staudt, N. Wali, J. Collins, S. Geyer, E. M. Busch-Nentwich, A. Galli, J. C. Smith, E. Robertson, D. J. Adams, W. J. Weninger, T. Mohun and M. Hemberger (2018). “Placentation defects are highly prevalent in embryonic lethal mouse mutants.” Nature 555(7697): 463–468.

Powers, S. K., L. L. Ji, A. N. Kavazis and M. J. Jackson (2011). “Reactive oxygen species: impact on skeletal muscle.” Compr Physiol 1(2): 941–969.

Prasasya, R., K. V. Grotheer, L. D. Siracusa and M. S. Bartolomei (2020). “Temple syndrome and Kagami-Ogata syndrome: clinical presentations, genotypes, models and mechanisms.” Hum Mol Genet 29(R1): R107–R116.

Riordan, J. D., V. W. Keng, B. R. Tschida, T. E. Scheetz, J. B. Bell, K. M. Podetz-Pedersen, C. D. Moser, N. G. Copeland, N. A. Jenkins, L. R. Roberts, D. A. Largaespada and A. J. Dupuy (2013). “Identification of rtl1, a retrotransposon-derived imprinted gene, as a novel driver of hepatocarcinogenesis.” PLoS Genet 9(4): e1003441.

Rodriguez-Martin, B., E. G. Alvarez, A. Baez-Ortega, J. Zamora, F. Supek, J. Demeulemeester, M. Santamarina, Y. S. Ju, J. Temes, D. Garcia-Souto, H. Detering, Y. Li, J. Rodriguez-Castro, A. Dueso-Barroso, A. L. Bruzos, S. C. Dentro, M. G. Blanco, G. Contino, D. Ardeljan, M. Tojo, N. D. Roberts, S. Zumalave, P. A. Edwards, J. Weischenfeldt, M. Puiggros, Z. Chong, K. Chen, E. A. Lee, J. A. Wala, K. M. Raine, A. Butler, S. M. Waszak, F. C. P. Navarro, S. E. Schumacher, J. Monlong, F. Maura, N. Bolli, G. Bourque, M. Gerstein, P. J. Park, D. C. Wedge, R. Beroukhim, D. Torrents, J. O. Korbel, I. Martincorena, R. C. Fitzgerald, P. Van Loo, H. H. Kazazian, K. H. Burns, P. S. V. W. Group, P. J. Campbell, J. M. C. Tubio and P. Consortium (2020). “Pan-cancer analysis of whole genomes identifies driver rearrangements promoted by LINE-1 retrotransposition.” Nat Genet 52(3): 306–319.

Rossant, J. and J. C. Cross (2001). “Placental development: lessons from mouse mutants.” Nat Rev Genet 2(7): 538–548.

Sala, L., S. Chandrasekhar and J. A. Vidigal (2020). “AGO unchained: Canonical and non-canonical roles of Argonaute proteins in mammals.” Front Biosci (Landmark Ed) 25(1): 1–42.

Sala, L., M. Kumar, M. Prajapat, S. Chandrasekhar, R. L. Cosby, G. La Rocca, T. S. Macfarlan, P. Awasthi, R. Chari, M. Kruhlak and J. A. Vidigal (2023). “AGO2 silences mobile transposons in the nucleus of quiescent cells.” Nat Struct Mol Biol 30(12): 1985–1995.

Segel, M., B. Lash, J. Song, A. Ladha, C. C. Liu, X. Jin, S. L. Mekhedov, R. K. Macrae, E. V. Koonin and F. Zhang (2021). “Mammalian retrovirus-like protein PEG10 packages its own mRNA and can be pseudotyped for mRNA delivery.” Science 373(6557): 882–889.

Seitz, H., N. Youngson, S. P. Lin, S. Dalbert, M. Paulsen, J. P. Bachellerie, A. C. Ferguson-Smith and J. Cavaille (2003). “Imprinted microRNA genes transcribed antisense to a reciprocally imprinted retrotransposon-like gene.” Nat Genet 34(3): 261–262.

Sekita, Y., H. Wagatsuma, K. Nakamura, R. Ono, M. Kagami, N. Wakisaka, T. Hino, R. Suzuki-Migishima, T. Kohda, A. Ogura, T. Ogata, M. Yokoyama, T. Kaneko-Ishino and F. Ishino (2008). “Role of retrotransposon-derived imprinted gene, Rtl1, in the feto-maternal interface of mouse placenta.” Nat Genet 40(2): 243–248.

Takahashi, M., M. Tamura, S. Sato and K. Kawakami (2018). “Mice doubly deficient in Six4 and Six5 show ventral body wall defects reproducing human omphalocele.” Dis Model Mech 11(10).

Toth, K. F., D. Pezic, E. Stuwe and A. Webster (2016). “The piRNA Pathway Guards the Germline Genome Against Transposable Elements.” Adv Exp Med Biol 886: 51–77.

Ulbricht, A., S. Gehlert, B. Leciejewski, T. Schiffer, W. Bloch and J. Hohfeld (2015). “Induction and adaptation of chaperone-assisted selective autophagy CASA in response to resistance exercise in human skeletal muscle.” Autophagy 11(3): 538–546.

van den Akker, E., M. Reijnen, J. Korving, A. Brouwer, F. Meijlink and J. Deschamps (1999). “Targeted inactivation of Hoxb8 affects survival of a spinal ganglion and causes aberrant limb reflexes.” Mech Dev 89(1-2): 103–114.

Vastenhouw, N. L. and R. H. Plasterk (2004). “RNAi protects the Caenorhabditis elegans germline against transposition.” Trends Genet 20(7): 314–319.

Watanabe, T., Y. Totoki, A. Toyoda, M. Kaneda, S. Kuramochi-Miyagawa, Y. Obata, H. Chiba, Y. Kohara, T. Kono, T. Nakano, M. A. Surani, Y. Sakaki and H. Sasaki (2008). “Endogenous siRNAs from naturally formed dsRNAs regulate transcripts in mouse oocytes.” Nature 453(7194): 539–543.

Wells, J. N. and C. Feschotte (2020). “A Field Guide to Eukaryotic Transposable Elements.” Annu Rev Genet 54: 539–561.

Wicker, T., F. Sabot, A. Hua-Van, J. L. Bennetzen, P. Capy, B. Chalhoub, A. Flavell, P. Leroy, M. Morgante, O. Panaud, E. Paux, P. SanMiguel and A. H. Schulman (2007). “A unified classification system for eukaryotic transposable elements.” Nat Rev Genet 8(12): 973–982.

Yekta, S., I. H. Shih and D. P. Bartel (2004). “MicroRNA-directed cleavage of HOXB8 mRNA.” Science 304(5670): 594–596.

Youngson, N. A., S. Kocialkowski, N. Peel and A. C. Ferguson-Smith (2005). “A small family of sushi-class retrotransposon-derived genes in mammals and their relation to genomic imprinting.” J Mol Evol 61(4): 481–490.

Zamore, P. D., T. Tuschl, P. A. Sharp and D. P. Bartel (2000). “RNAi: double-stranded RNA directs the ATP-dependent cleavage of mRNA at 21 to 23 nucleotide intervals.” Cell 101(1): 25–33.

